# A new corrector rescues F508del-CFTR folding through stabilization of the TMD1-NBD1 linker

**DOI:** 10.1101/2025.03.10.640553

**Authors:** Peter van der Sluijs, Olga Papaioannou, Azib Hagos, Aya M. Saleh, Timothy Hodges, Corina Balut, Matthew Green, Sara Alani, Yihong Fan, Kazumi Ebine, Michael Schrimpf, David Hardee, Huan-Qiu Li, Henry Korman, Laura Tade, Guus van Zadelhoff, Ashvani Singh, Chris Tse, Ineke Braakman

## Abstract

While Trikafta has turned many CF patients into people that lead relatively healthy lives, additional modulators are still needed. We developed an acylsulfonamide-type C2 corrector X307810 that is potent and shows a distinctive mode of action in rescuing F508del-CFTR from degradation at the endoplasmic reticulum. We tracked the de-novo folding and assembly of each CFTR domain with a radio-active pulse chase - limited proteolysis folding assay and determined when and where X307810 works on the newly synthesized protein. X307810 rescued the Golgi form of F508del-CFTR to 80% of wild-type values. To our knowledge, such a dramatic increase has never been attained with a single corrector. Zooming in on X307810 mode of action revealed that the corrector changed protease resistance of the linker between TMD1 and NBD1 during late stages of CFTR folding. X307810 is the first corrector that added a unique new fragment to the typical limited-proteolysis pattern. Other C2 correctors do not change fragments, but portray an increase in fragment quantity characteristic of domain assembly in wild-type and corrected F508del-CFTR. X307810 instead extended protection of TMD1 from intracellular loop 2 to the unstructured regulatory insertion of NBD1. These results provide new and important insights into the impact of acylsulfonamide correctors on CFTR structure and function.

**SIGNIFICANCE STATEMENT:** CFTR is an anion channel, mutations in which cause cystic fibrosis (CF). Corrector compounds allow misfolded missense CFTR mutants to escape from degradation in the endoplasmic reticulum, which is the basis for CF therapy, in combination with a potentiator. How correctors improve folding is incompletely understood. We here report on X307810 having a novel mode of action on post-translational domain assembly of newly synthesized CFTR. Our in-cell folding assay shows that X307810 stabilizes the linker between the first transmembrane domain (TMD1) and the first nucleotide binding domain (NBD1) into the Regulatory Insertion; it strengthens the TMD1-NBD1-ICL4 interface. The action of X307810 boosts egress of F508del-CFTR from the ER to an unprecedented 80% for a single modulator.

## INTRODUCTION

CFTR (Cystic Fibrosis Transmembrane Conductance Regulator) is an anion channel that mediates chloride and bicarbonate export in a passive but regulated mode at the apical plasma membrane of epithelial cells in lung and intestine (1). It belongs to the ABCC subfamily of the superfamily of ATP-binding cassette (ABC) transporters and consists of two transmembrane domains (TMDs) and two nucleotide-binding domains (NBDs), both characteristic of ABC transporters (2). CFTR has a unique regulatory, intrinsically disordered region (R) that connects the two halves of the protein (3). Like other ABCC proteins, CFTR features an N-terminal cytoplasmic lasso motif, which contributes to stability and interactions with other proteins (4) but whose role in CFTR biology is not well understood (2). CFTR function is regulated by more than 10 protein kinase A-dependent phosphorylation events in the R domain in addition to ATP binding and hydrolysis (5).

Over 4,000 mutations have been identified in CFTR (6), of which >700 were shown to cause CF (7). The most common CF-causing mutation is the deletion of amino acid phenylalanine at position 508 (F508del). The resulting severely dysregulated chloride and water homeostasis give rise to thick mucus that can clog secretory organs and airways including lungs and pancreas (8). Missense mutations can cause reduced levels or complete absence of functional channels on the cell surface (9), due to misfolding in the endoplasmic reticulum (ER) and ensuing degradation by the proteasome and autophagy (10). Through a variety of well-characterized cellular expression systems, the first principles of CFTR proteostasis, its maturation, and regulation at the plasma membrane have emerged (11, 12). Since missense mutations still permit synthesis of a non-functional full-length protein, albeit (oftentimes) with reduced half-life, these can be targeted in compound screens for CF therapies (10).

Over the years, improvements in symptomatic treatments advanced the median survival age of CF patients from 32 years in 1990 to 50 years in 2012 (13). In 2009, the first clinical modulators that act directly on the CFTR protein became available, which proved a watershed in CF treatment and led to a stellar increase in life expectancy of many CFTR patients (14). Current small-molecule modulators can be divided roughly into two classes, namely potentiators that improve gating at the cell surface and correctors that enhance transport of misfolded CFTR to the cell surface. Current standard of care consists of a triple combination consisting of potentiator VX770 (ivacaftor), which interacts with residues in transmembrane helices TM4, TM5, and TM8, corrector VX661 (tezacaftor), which binds TM2, TM3, and TM6, and a third corrector VX445 (elexacaftor), which binds TM2, TM11, and the Lasso (15, 16). How correctors work in the cell is incompletely understood and CFTR structures that include modulators have been obtained through incubation with folded recombinant CFTR protein, while correctors in the cell act on CFTR molecules that are in the process of vectorial folding but have not yet reached the functional form (16, 17). Progressive insight uncovered that VX445, which originally was typecast as a novel type of corrector, also has potentiator activity (18). A binary classification of drugs for CFTR rescue is therefore likely too simplistic, especially since medicinal-chemistry campaigns in combination with artificial intelligence and machine learning may yield new therapeutic compounds with unanticipated modes of action on CFTR (19–21).

We have developed an assay to track the folding of newly synthesized proteins in cells that allows analyzing the effects of small-molecule modulators on CFTR conformation during the in-vivo folding process. We have found that CFTR folding occurs in 2 discrete, partially overlapping stages, representing a modular pathway of individual domain folding followed by assembly of the domains into the final functional structure (22).

The division of the folding pathway in 2 distinct overlapping sequential stages provides a Janus face to pharmacological intervention of CF. The challenge of this arrangement is that a mutation not only can destroy the domain in which the mutation is situated, but the mutation can also block entry into or completion of the second stage. An example of which is the most abundant disease-causing mutation F508del, where deletion of F508 precludes co-translational folding of NBD1 as well as posttranslational assembly with the other 3 domains. That not all domain interfaces are critical for assembly is shown by rescue of F508del-CFTR folding by VX445, VX661, and VX770 to near-wild-type levels (22, 23). The 3 other domains stabilize the defective domain upon assembly, leading to functional improvement. F508del-NBD1 thermodynamic instability and folding, however are *not* improved by these drugs; nor does low temperature rescue F508del-NBD1 folding (10, 22).

Although the available triple modulators therapy has revolutionized the treatment of CF, still at least 10% of CF patients do not benefit from Trikafta, either because their CFTR mutations are refractory to treatment, or because of the complexity of the disease. Additional factors such as high plasma-protein binding (14), interactions with cytochrome P450 (24), drug-drug interactions (25), adverse effects even though rare (26), and the price legitimate the search for a broader range of correctors. This would allow treatment to be tailored to more specific mutations, thus covering a larger proportion of the population with CF.

We here report the development and mode-of-action analysis of a new class of acylsulfonamide-type corrector compounds. Two of these, X307810 and X323022, individually rescued misfolded F508del-CFTR from the ER to the same extent as VX445 plus VX661. X307810 acts additively with the C1-type corrector X281602 (27) and together these stimulate export from the ER more efficiently than VX445 and VX661. In-cell folding assays uncovered a new mode of action on CFTR folding, in which X307810 stabilizes the linker between TMD1 and NBD1 of F508del and wild-type CFTR during (induced) posttranslational assembly of the domains. Improved tethering of (misfolded) F508del NBD1 onto TMD1 thereby relieves the block on posttranslational domain assembly and concurrently facilitates transport to the Golgi complex and beyond.

## RESULTS

### Discovery of substituted cyclopropane CFTR correctors

To develop novel CFTR correctors, we began by investigating the chemical matter related to a family of known C2 corrector molecules exemplified by compound **1** (Figure 1A). (REF: US 2019/0077784). The goal of this effort was to enhance C2 corrector activity as measured by TECC and Cell Surface Expression (CSE) while also achieving a desirable *in-vivo* pharmacokinetic profile (the latter is beyond the scope of this manuscript). Pursuant to this objective, we prospected the previously unassessed chemical space around the cyclopropyl linker connecting the left-hand aryl region to the right-hand acyl sulfonamide moiety.

**Figure 1:**
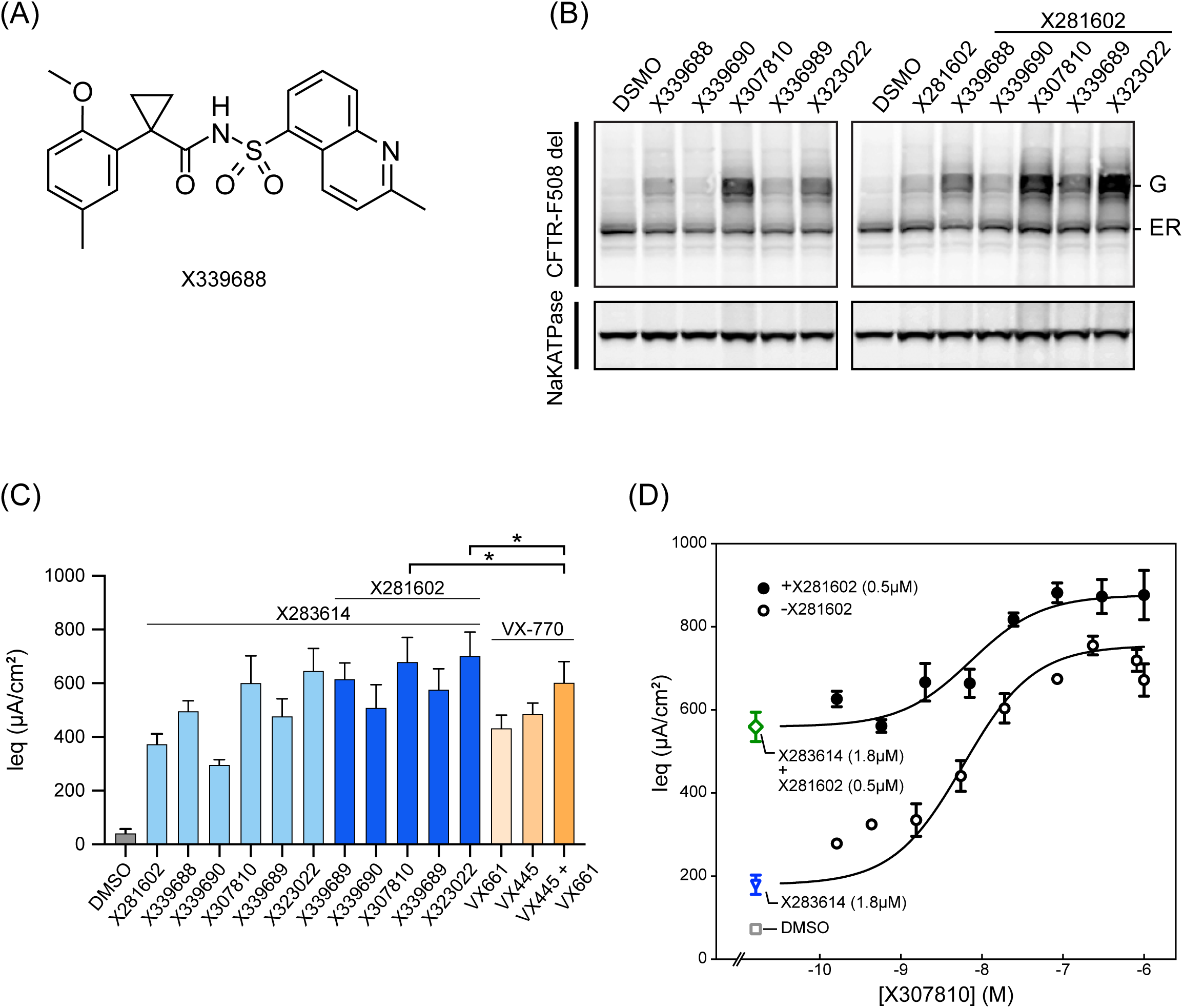
Synthesis and pharmacological characterization of novel modulators. **A)** Structural formula of starting compound X339688 (compound 1) for synthesis of modulators with enhanced CFTR corrector activity. **B)** CFBE410-cells stably expressing CFTR508del were treated overnight with 3 μM of indicated compounds and analyzed by Western blot using antibodies against CFTR and Na^+^/K^+^ ATPase. Positions of oligomannose ER form (ER) and complex-glycosylated Golgi form (G) are indicated. **C)** Bar graph showing mean current values obtained in TECC assays with HBE cells homozygous for F508del-CFTR. Data are means ± SD of at least 5 experiments, *denotes p < 0.05 (one-tailed t test). **D)** Dose-response (current) curve of X307810 in the presence of the potentiator X283614 (1.8 µM) using primary HBE cells as in B. The EC_50_ for X307810 in the post-screen TECC assays was 5.9 nM, data are means ± SD of at least 3 experiments.

Early efforts in functionalizing the cyclopropane afforded methylene methyl ether enantiomers **2** and **3** (Table 1). While compound **2** maintained a similar functional activity (TECC EC_50_) relative to unsubstituted **1**, compound **3** showed a 10-fold decrease in potency. Switching to the larger and more lipophilic phenyl group afforded analogs **4** and **5** with enhanced TECC EC_50_’s of 0.99 nM and 11.3 nM, respectively. Notably, these results further demonstrated a significant eudysmic ratio for substitution at this position and a dramatically higher maximum cell surface expression (3360%) for compound **4** relative to parent compound **1**.

**Table 1:** CFTR activity for select substituted cyclopropane molecules.

| compound | R | Stereo (c<sup>1</sup>,c<sup>2</sup>) | TECC EC<sub>50</sub> (nM)<sup>a, c</sup> | TECC Max (%)<sup>b, c</sup> | CSE-HRP EC<sub>50</sub> (μM)<sup>d</sup> | CSE-HRP Max (%)<sup>d</sup> |
| --- | --- | --- | --- | --- | --- | --- |
| **X339688 (1)** | H | N/A | 27.5 | 119 | 1.18 | 1630 |
| **(2)** | CH<sub>2</sub>OCH<sub>3</sub> | (S,S) | 26 | 96 | 0.956 | 1000 |
| **(3)** | CH<sub>2</sub>OCH<sub>3</sub> | (R,R) | 281 | 96 | 2.05 | 715 |
| **X307810 (4)** | Ph | (S,R) | 0.99 | 127 | 0.994 | 3360 |
| **X339689 (5)** | Ph | (R,S) | 11.3 | 107 | 0.497 | 1190 |
| **X339690 (6)** | 2-Py | (S,S) | 449 | 97 | 2.8 | 1190 |
| **(7)** | 6-Me-2-Py | (S,S) | 22.7 | 130 | 0.99 | 1990 |
| **X323022 (8)** | 6-OEt-2-Py | (S,S) | 1.67 | 149 | 0.339 | 3120 |

<sup>a</sup>Dose-response curves and EC<sub>50</sub> estimates for compounds **1-8** were done in the presence of 0.15 μM X281605 (C1 corrector) + 0.45 μM X283649 (potentiator).
<sup>b</sup>TECC assay was run using 1 μM X289990 (C2 corrector) + 0.15 μM X281605 + 0.45 μM X283649 as a positive control. Reported values are relative to this positive control.
<sup>c</sup>Mean of 2 or more independent experiments.
<sup>d</sup>Results from a single experiment.

Appending the phenyl group to the cyclopropyl linker improved the *in vitro* corrector activity but also increased the lipophilicity of the molecule. To reduce lipophilicity, polar heterocyclic cyclopropane substituents were examined as phenyl replacements. As such, a nitrogen atom was introduced into the cyclopropane substituent to afford pyridyl compound **6**, which resulted in a dramatic loss of TECC potency. We hypothesized that the basicity of the pyridyl group was responsible for the reduced activity. To mask this basicity, methyl pyridine **7** was synthesized and exhibited an almost 20-fold improvement in TECC EC_50_ relative to compound **6**. Further enhancements of C2 corrector activity were achieved by substituting the pyridine ring with ether groups as exemplified by ethoxy pyridine **8**, that featured a TECC EC_50_ of 1.67 nM, a max cell surface expression of 3120%, and compared favorably to the activity of phenyl substituted **4**.

### Effect of novel AbbVie (ABBV) correctors on F508del-CFTR maturation in CFBE41o-cells

We incubated CFBE41o-cells expressing F508del-CFTR overnight with 3 μM of the corrector compounds without and with the AbbVie C1 corrector X281602 to determine their effect on steady-state levels of F508del-CFTR. Compared to the other compounds, X307810 and X323022 greatly improved F508del-CFTR maturation as indicated by the accrual in intensity of the complex-glycosylated Golgi (G) band (Figure 1B). CFTR folding and transport are efficient and robust processes, which we (10, 22, 28, 29) and others (30–32) have shown to be remarkably similar between such diverse cells as CFBE, HeLa, and FRT. We therefore performed a more quantitative evaluation of the AbbVie corrector compounds in HEK293T cells expressing F508del-CFTR (Figure S2A). Except for X339690, all compounds enhanced the amount of F508del-CFTR in the Golgi complex (Figure S2B). We found that X307810 and X323022 individually performed better than VX445 or VX661, but were equally effective to VX445 and VX661 in combination (Figure S2B). Combining X281602 with any of the new corrector compounds increased the amount of F508del-CFTR in the Golgi complex compared with X281602 alone (Figure S2B). Also, X281602 increased the effect of each corrector compound on export of F508del-CFTR from the ER, although this did not reach significance for X281602 + X323022 (p = 0.06; Figure S2C). The absence of additivity of X281602 and X323022 is explained by the efficacy of X323022, which by itself already reaches near wild-type levels of ER-to-Golgi transport. A direct comparison with the Vertex correctors showed that X281602 + X307810 or X281602 + X323022 improved F508del-CFTR corrector activity more than VX445 + VX661 (p < 0.05).

To evaluate whether the unusually efficient rescue of F508del-CFTR by X307810 or X323022 from the ER could be ascribed to dual corrector activity, we used additional assays for characterizing C1 corrector activity. As we know, true C1 correctors target TMD1 and stabilize the domain against degradation (33, 34). We therefore incubated HEK293T cells expressing TMD1 with X307810 during starvation and pulse chase. After pulse labeling, cells were chased for up to 4 h in the presence of cycloheximide and TMD1 was immunoprecipitated from the cell lysates (Figure S3A). Turnover of TMD1 was not changed by the presence of X307810 versus DMSO controls, which is inconsistent with compounds having C1 activity. We then used the radiolabeling-limited-proteolysis *in cell* folding assay to determine the corrector compound’s activity during co-translational folding of F508del-CFTR. HEK293T cells expressing F508del-CFTR were incubated with 3 µM of X307810 or X281602 during starvation and pulse labeling; cell lysates were subjected to limited proteolysis and immunoprecipitation with TMD1 antibodies (Figure S3B). We found an increase in the TMD1-derived T1aa fragment in the presence of X281602 but not X307810. T1aa is a TMD1 fragment consisting of residues 36-256 (35) that features a slightly lower electrophoretic mobility than T1a (residues 50-256) (22) because the region around residue 50 of the cytoplasmic N-terminus has folded into a structure that protects it from digestion by ProtK. The gain in T1aa is a characteristic signature of class-1 correctors, which stabilize TMD1 during its co-translational folding by enhancing early packing of the transmembrane helices as we established for VX809 (35) and posenacaftor (not shown). The new correctors X307810 and X323022 thus have a different mode of action than C1 correctors.

### Functional characterization of corrector compounds in primary HBE cells

To determine the pharmacological efficacy of ABBV correctors, HBE cells derived from CF patients homozygous for the F508del mutation were incubated with the compounds 24 h prior to electrophysiological measurements using a TECC functional assay (Figure 1C). CFTR-mediated equivalent current (I_eq_) was measured for correctors in the presence of potentiator X283614. In line with the Western blot results (Figure 1B and Figure S2), X307810 and X323022 exhibited the highest activity among ABBV correctors tested. Additionally, both correctors showed slightly higher efficacy compared to VX-445 in the presence of potentiator VX-770 (p < 0.05). A dose-response curve for X307810 in the presence of potentiator X283614 yielded an EC_50_ of 5.9 nM, a value slightly higher than in the initial screens (Table I). We did not measure a further increase in Max I_eq_ values when we added the C1 corrector X281602 at 0.5 μM (Figure 1D), however, there was a marked increase in the baseline activity indicating the activity of the C1 corrector. Taken together, the biochemical and functional results signify that X307810 and X323022 are potent corrector compounds with improved *in vitro* efficacy compared to VX-445.

### X307810 and X323022 rescue F508del-CFTR via a distinct mode of action

We next used the in cell folding assay to evaluate the corrector compounds on newly synthesized F508del-CFTR. HEK293T cells transiently expressing F508del-CFTR were incubated during starvation and pulse chased with 3 μM of the corrector compounds. After 2 h of chase, the amount of the core-glycosylated ER form of CFTR decreased, and only a tiny amount of F508del-CFTR was exported out of the ER as shown by the very faint Golgi band (Figure 2A, top panel). All correctors enhanced export of F508del-CFTR to the Golgi complex, but X307810 and X323022 did so most efficiently in accord with the steady-state analysis by Western blot (Figure 1B). Both correctors surpassed VX-445 (Figure 2A, top panel) in comparison. To establish F508del-CFTR conformation, we immunoprecipitated TMD1 fragments from proteolyzed lysates of corrector-treated cells at 2-h chase, which yielded the well-established T1d, T1e, and T1f bands. These fragments constitute the signature of post-translationally folded, domain-assembled CFTR (22) and are not seen in F508del-CFTR controls nor from cells expressing wild-type or F508del-CFTR that were lysed immediately after the pulse, where folding had not proceeded beyond domain folding (22). Larger amounts of T1d-f fragments were generated from F508del-CFTR treated with X307810 and X323022, due to increased folding, increased domain assembly and enhanced packing. This reflects protection of ICL2 and the complete cytoplasmic N-terminus, in accord with the wrapping of the N terminus around the TMs of TMD1 and TMD2 which correlates with more efficient rescue of full-length protein from the ER. None of the correctors rescued folding of NBD1 as evidenced by the absence of the characteristic wild-type N1a fragment, similar to the effects of VX-809 and VX-445 (22).

**Figure 2:**
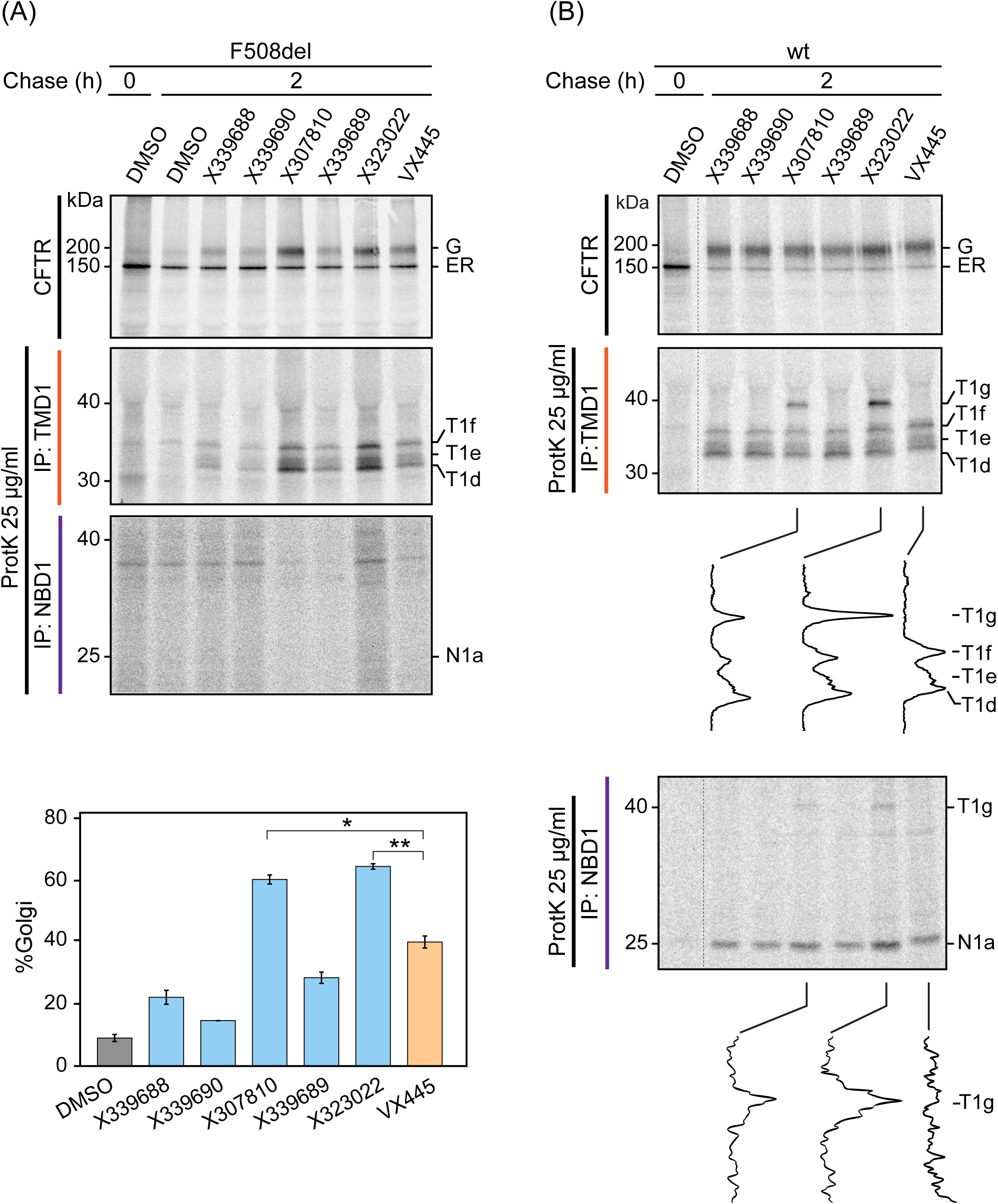
Effect of modulators on de-novo folding. **A)** HEK293T cells expressing wild-type or F508del-CFTR were treated with modulator compounds at 3 μM final concentration during starvation, radiolabeling, and chase. Cells were pulse labeled for 15 minutes with ^35^S-Met/^35^S-Cys and chased for 2 h or not (0 h) before detergent lysis. CFTR was immunoprecipitated using MrPink and immunoprecipitates were resolved by 7.5% SDS-PAGE. Remaining lysates were subjected to limited proteolysis with 25 µg/mL ProtK. Protease-resistant fragments were immunoprecipitated with E1-22 (TMD1) or MrPink (NBD1), and resolved by 12% SDS-PAGE. T1d-f are TMD1-specific protease-resistant fragments; N1a is a protease-resistant fragment derived from NBD1. Bar graph with % CFTR transported to the Golgi complex (G*100/(G+ER)) in the presence of compounds. Data are means ± SD of 3 experiments. * indicates p < 0.005 and ** p < 0.001 in one-tailed t tests). **B)** HEK293T cells expressing wild-type (wt) CFTR received modulators and were processed as in **2A)**. ER and G denote positions of oligomannose and complex-glycosylated forms, respectively. Band intensities in selected lanes of the T1d - T1g region were scanned to highlight the presence of the T1g fragment in the wild-type and F508del-CFTR proteolytic digests immunoprecipitated with TMD1 and NBD1 antibodies. Note the T1g fragment in CFTR digests from X307810-treated and X323022-treated cells.

We then analyzed the effect of correctors on folding of wild-type CFTR. Differences between effectors were less pronounced than for F508del-CFTR, likely because wild-type CFTR is already efficiently exported from the ER to the Golgi and beyond. Limited proteolysis followed by immunoprecipitation with TMD1 and NBD1 antibodies yielded the T1d-f signature bands (22) for the TMD1 immunoprecipitates and in addition a 40-kDa band, which we coined T1g (Figure 2B) to denote its origin as TMD1-derived. The T1g fragment generated with limited proteolysis from wild-type CFTR is a stable proteolytic intermediate that has not been detected before, neither with nor without small modulator compounds. The 40-kDa fragment was immunoprecipitated with the MrPink NBD1 antibody as well (Figure 2B, bottom panel), although less efficiently than with the TMD1 antibody. T1g therefore contains TMD1 as well as part of NBD1.

### The 40-kDa folding intermediate of F508del-CFTR is less stable than that of wild-type CFTR

Interestingly, T1g was not detected upon limited proteolysis of F508del-CFTR und under identical conditions er identical conditions (Figure 2A). It is known that a floppy F508del NBD1 is more accessible and susceptible to ProtK than correctly folded wild-type NBD1. The standard conditions in the limited-proteolysis assay therefore may prohibit detection of the T1g fragment from F508del NBD1. We titrated the ProtK concentration over 4 orders of magnitude to capture a possibly transient intermediate. F508del-CFTR cells were radiolabeled for 15 min and chased for 2 h in the presence or absence of X307810 (Figure S4A). Decreasing ProtK concentration 10-fold revealed a strong band of ∼40 kDa with similar mobility as T1g derived from wild-type CFTR (Figure S4B), after immunoprecipitation with the E1-22 antibody. The same band was also generated when we used the standard ProtK concentration but reduced the time of proteolysis (Figure S4C) from 15 to 1 min. We then evaluated whether the novel corrector compounds had the same mode of action on F508del-CFTR, by comparing effects of X307810, X323022, and the less active X339688 (Figure 1B and S2) in folding assays with 10 and 100-fold dilution of ProtK (Figure 3A). Limited proteolysis with 10-fold ProtK dilution yielded the T1g band from F508del-CFTR treated with any of the three compounds (Figure 3B). Less of the T1g was generated from lysates of X339688-treated cells in accord with more efficient rescue of full-length F508del-CFTR by X307810 and X323022 (Figure 3A). The low protease concentration did not yield an N1a fragment from F508del-CFTR, confirming that F508del NBD1 remained misfolded upon corrector treatment (Figure 3C). Since the effects of X307810 and X323022 on F508del-CFTR were largely indistinguishable, we continued characterization of X307810 only.

**Figure 3:**
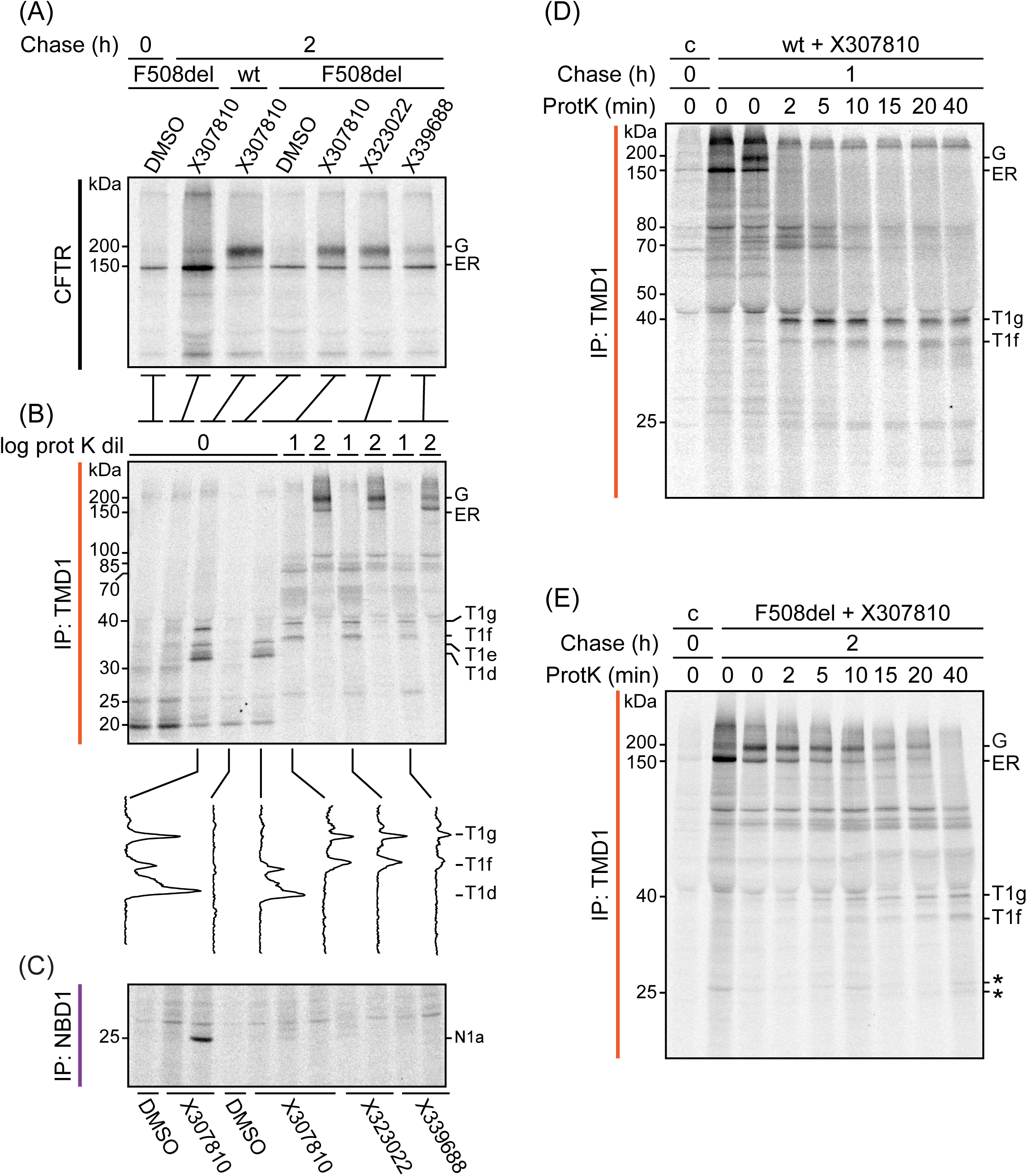
Modulator compounds stabilize the TMD1-NBD1 linker in F508del-CFTR. **A)** HEK293T cells expressing wild-type (wt) or F508del-CFTR were treated with corrector compounds labeled and chased as in Figure 2A. CFTR was immunoprecipitated using MrPink and immunoprecipitates were resolved by 7.5% SDS-PAGE. **B)** Remaining lysates were subjected to limited proteolysis with 25 (0), 2.5 (1) or 0.25 µg/mL (2) ProtK and protease-resistant fragments were immunoprecipitated with E1-22 antibody and resolved by 12% SDS-PAGE. Band intensities in selected lanes of the T1d - T1g region were scanned to highlight the presence of the T1g proteolytic fragment from wild-type and F508del-CFTR. **(C)** As B but digested lysates were immunoprecipitated with MrPink antibody against NBD1; note the absence of N1a in all F508del-NBD1-derived lanes with or without compounds. (**D**) HEK293T cells expressing wild-type (wt) CFTR or **(E)** F508del-CFTR were incubated with X307810, radiolabeled and chased as in Figure 2A. Full-length CFTR was immunoprecipitated with E1-22 and MrPink (Figure S5A,B) antibodies and analyzed on 7.5% SDS-PAA gels. Remaining lysates were digested with 5 μg/mL ProtK for wild-type CFTR, or 0.5 μg/mL for F508del-CFTR for different periods of time. Proteolytic fragments were immunoprecipitated with E1-22 against TMD1 and MrPink against NBD1 (Figure S5C,D), and analyzed by 12% SDS-PAGE. Asterisks denote background bands.

To establish precursor-product relationships for T1g fragment formation from full-length wild-type CFTR in cells treated with X307810, we performed limited-proteolysis time-course experiments. The dose of ProtK was reduced to 5 μg/ml to maximize chances of detecting transient intermediate proteolytic TMD1 fragments. Cells were radioactively labeled with ^35^S methionine and cysteine for 15 min, chased for 2 h and lysed. A small fraction was saved for analysis of full-length protein (Figure S5A); the remaining lysate was subjected to limited proteolysis with ProtK for up to 45 min and proteolytic fragments were immunoprecipitated with E1-22 (Figure 3D) and MrPink antibodies (Figure S5C). Within 2 min, full-length CFTR was cleaved to a fragment of 72 kDa, which was converted within 10 min to lower-molecular-weight species. T1g and T1f were generated in 2 min as well but were more stable and started to decline gradually after 15 min. Immunoprecipitations with MrPink confirmed that NBD1, in addition to TMD1, is part of the fragment with the 72-kDa band (Figure S5C). This apparent molecular weight suggested that the cleavage occurred in the intrinsically disordered R region, generating the two hemi-transporter halves of CFTR. We also subjected lysates of X307810 -treated F508del-CFTR-expressing cells to a time course with an attenuated concentration (0.5 μg/ml, 10-fold less than for wild-type CFTR) of ProtK (Figure 3E). In contrast to wild-type CFTR in Figure 3D, small amounts of T1g were immunoprecipitated with the TMD1 antibody at the first time point, and the fragment was relatively stable, while T1f lagged behind the formation of T1g. Cleavage of full-length F508del-CFTR into and within the half transporter was protracted most likely because of the low ProtK concentration (Figure 3E). Immunoprecipitation with the MrPink antibody did not yield detectable T1g fragment nor the 25-kDa N1a fragment (Figure S5D) characteristic of properly folded NBD1 (22).

### TMD2 and NBD2 are efficiently rescued by X307810

De-novo folding of CFTR occurs in sequential but overlapping stages of co-translational domain folding and posttranslational assembly of domains into a functional molecule. The in-vivo folding assay allows interrogation of these stages separately through immunoprecipitation with domain-specific antibodies. This unveils landmark banding patterns by SDS-PAGE that are characteristic of early and late folding stages. We extended the analysis of the effect of X307810 on CFTR folding to TMD2 and NBD2 constituting the C-terminal half of (F508del) CFTR in comparison with VX-445 and VX-661. At the end of the 15 min radiolabeling, similar amounts of the core glycosylated (ER) form of full-length wild-type and F508del-CFTR were immunoprecipitated in the presence or absence of each compound (Figure 4A, left panel). Limited-proteolysis fragments for TMD1 were similar in all conditions and the N1a fragment was only seen for wildt NBD1. Slightly lower amounts of the early TMD2 fragment T2b were detected for F508 del-CFTR, and this was not improved by any of the compounds. Small amounts of the NBD2 limited-proteolysis fragment N2a were detected for wild-type but not F508del-CFTR, irrespective of the presence of compounds, consistent with a small fraction of wild-type CFTR molecules whose synthesis and folding had already been completed during the pulse-labeling period (Figure 4B, left panels).

**Figure 4:**
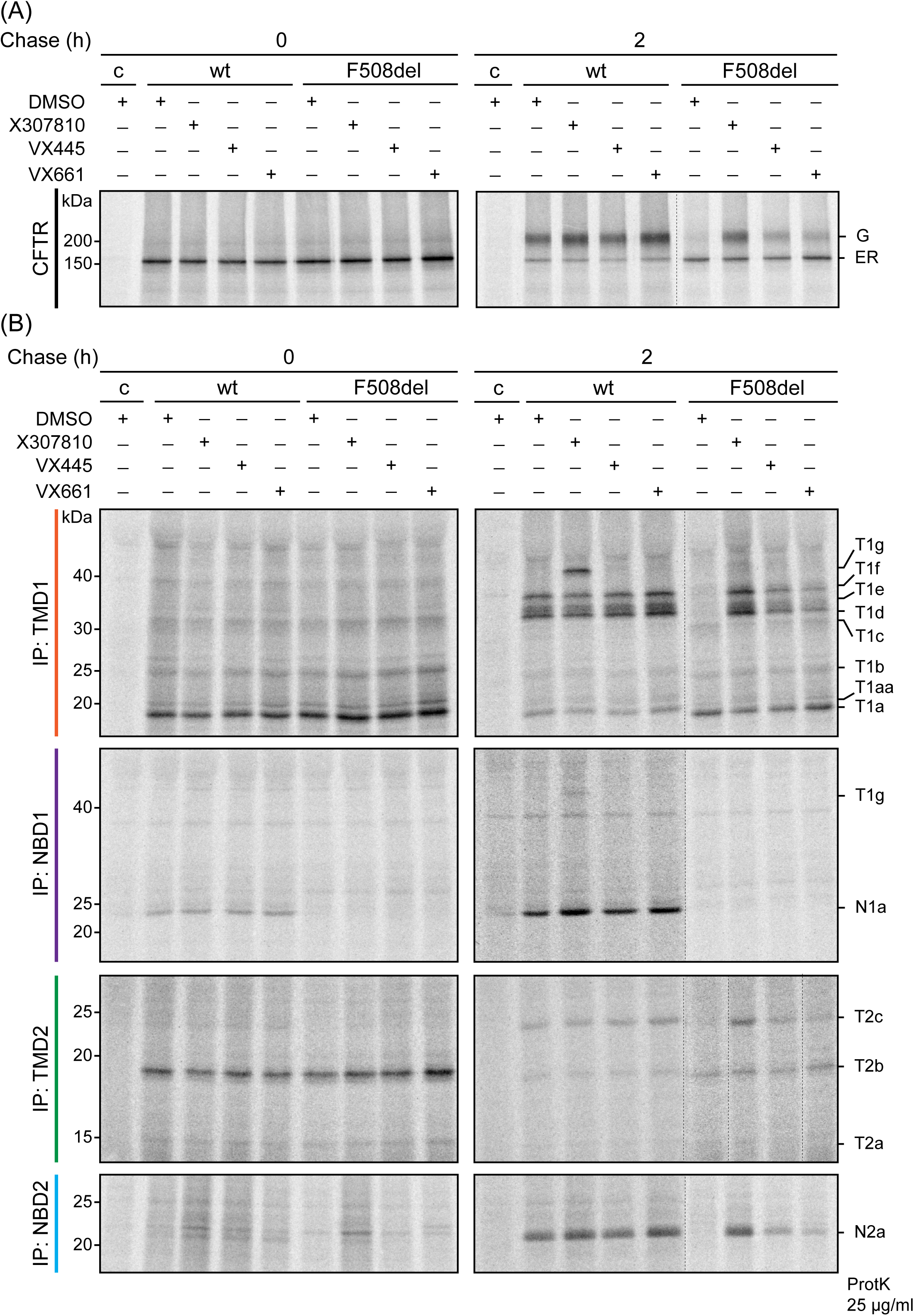
Corrector compound X307810 rescues domain assembly of F508del-CFTR. **(A)** HEK293T cells expressing wild-type (wt) or F508del-CFTR were incubated with corrector compounds, labeled and chased as in Figure 2A. CFTR was immunoprecipitated using MrPink and immunoprecipitates were resolved by 7.5% SDS-PAGE. **B)** Remaining lysates were subjected to limited proteolysis with 25 µg/mL ProtK and protease-resistant fragments were immunoprecipitated with E1-22 (TMD1) or MrPink (NBD1) and resolved by 12% SDS-PAGE. wild-type, wild-type CFTR; T1a-T1f are TMD1-specific protease-resistant fragments; T2a-c are TMD2-specific protease resistant fragments; N1a and N2a are protease-resistant fragments specific for NBD1 and NBD2, respectively.

After 2 h of chase, 82% of wild-type had received Golgi glycan modifications, and less than 10% of F508del-CFTR. In the presence of X307810, this value increased to 65% for F508del-CFTR, versus 37% and 40% for VX-445 and VX-661-treated F508del-CFTR, respectively (Figure 4A, right panel). Consistent with rescue of full-length F508del-CFTR by the modulators, limited proteolysis and immunoprecipitation with TMD1 antibody revealed an increase in T1d-f fragments characteristic of domain assembly by 21, 6, and 7-fold (compared to non-treated F508del-CFTR) for X307810, VX445, and VX-661, respectively. The increase of T1d-f for wild-type was lower in the presence of X307810, because precursor T1g was more protease resistant and therefore yielded less of the smaller T1def fragments (Figure 4B, right panels).

The T1g intermediate was most efficiently detected for X307810-treated cells, in both the wild-type TMD1 and NBD1 immunoprecipitations. Because the limited-proteolysis experiments were done with 25 μg/ml ProtK, the T1g band was not seen for corrected F508del-CFTR. A strong T1g signal was also detected in the NBD1 immunoprecipitate of proteolysate from X307810-treated wild-type cells, in accord with the TMD1 immunoprecipitation, but not for the NBD1 immunoprecipitation of F508del-CFTR. As expected from the domain-assembly-indicating fragments T1d-f from TMD1, X307810 also provided rescue of the domains in the C-terminal half of F508del-CFTR: the late TMD2 proteolytic fragment T2c was nearly undetectable from control F508del-CFTR, and increased 10, 2, and 3-fold for X307810, VX445, and VX-661-treated cells, respectively. The N2a proteolytic fragment of the NBD2 domain was rescued 11-fold by X307810, and 5 and 3-fold by VX-445, and VX661, respectively (Figure 4B, right panels).

### X307810 protects the TMD1-NBD1 linker and the Lasso during folding

Finally, understanding how the new corrector compounds affect the folding pathway of de-novo synthesized CFTR requires determination where the T1g proteolytic fragment maps on the CFTR sequence. Unfortunately, mass spectrometry does not yet have the requisite sensitivity to detect minute amounts of proteolytic fragments from the multispan membrane protein CFTR and would not offer the temporal resolution of folding stages during membrane-protein biogenesis. We therefore capitalized on results from previous work in which we developed strategies to determine the N- and C-terminal boundaries of the proteolytic fragments generated by ProtK from TMD1 (22). The essence is the use of N terminally truncated CFTR mutants in conjunction with immunoprecipitation of proteolytic fragments with CFTR antibodies that recognize spatially separated epitopes on CFTR (Figure 5A). Since T1g is larger than T1f, it likely has a longer C-terminus than T1f, and therefore may be detectable by antibodies with epitopes more C-terminal to T1f (Figure 5A). The N terminus of T1g likely coincides with that of T1f, which is the start methionine (Figure 5A), the most N-terminal amino acid of CFTR. Accordingly, short N-terminal truncations upstream of the N terminus of the T1d fragment (which is at residue 36 (Figure 5A)) shorten the T1f fragment, but have no effect on the mobility of the T1d fragment by SDS-PAGE (22).

**Figure 5:**
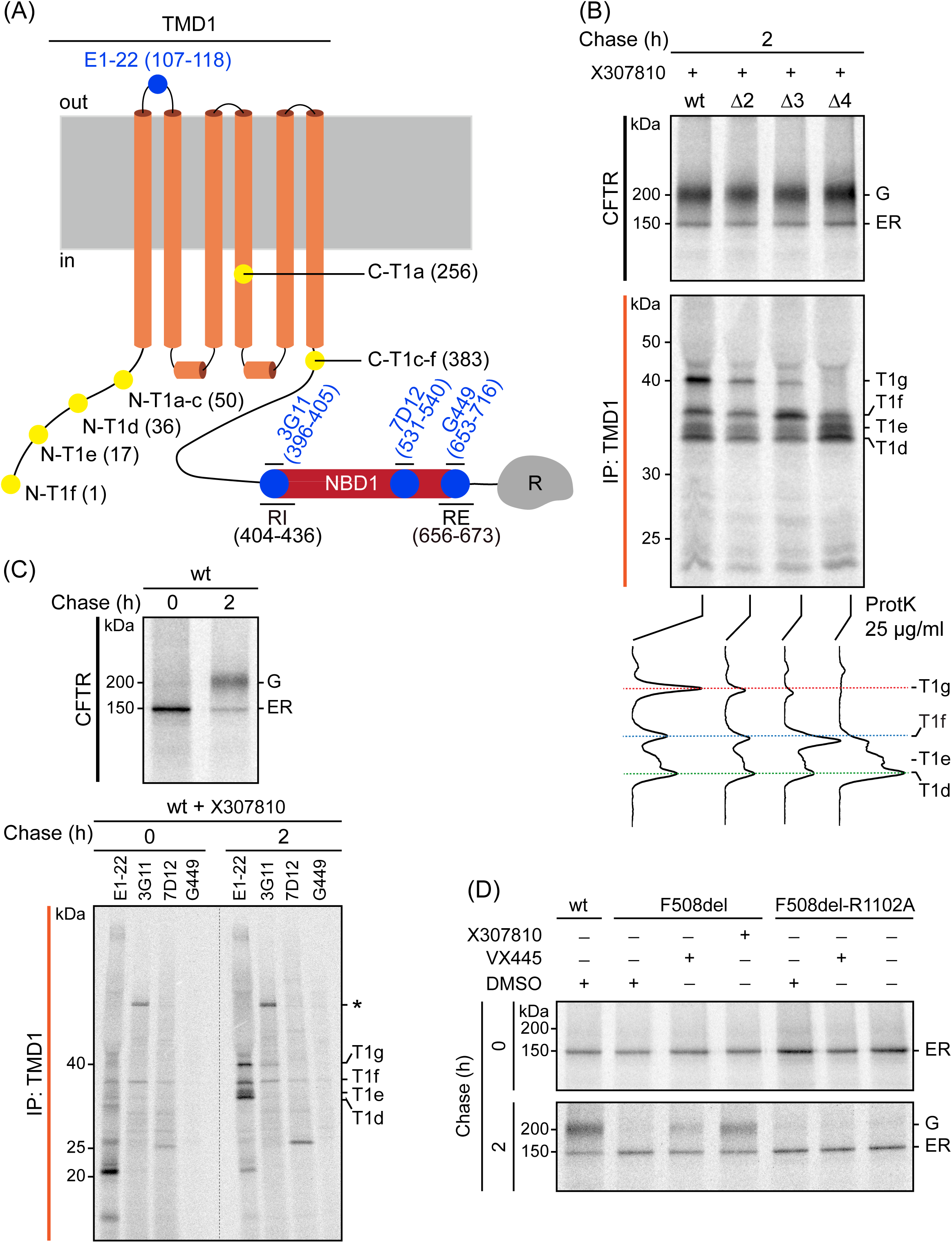
X307810 stabilizes the linker between TMD1 and RI of NBD1. **A)** Schematic with boundaries of TMD1 proteolytic fragments (filled yellow circles) and antibody epitopes (filled blue circles). **B)** HEK293T cells expressing N-terminally truncated wild-type (wt) CFTR variants were incubated with X307810, labeled, chased and lysed as in Figure 2A. CFTR was immunoprecipitated using MrPink and immunoprecipitates were resolved by 7.5% SDS-PAGE (top panel). Remaining lysates were subjected to limited proteolysis with 25 µg/mL ProtK and protease-resistant fragments were immunoprecipitated with E1-22 and resolved by 12% SDS-PAGE. Note the faster migrating T1f and T1g bands when deleting 2 or 3 amino acids from the N-terminus, while T1d and T1e remained invariant. **C)** HEK293T cells expressing wild-type (wt) CFTR were incubated with corrector compounds, labeled, chased and lysed as in Figure 2A. CFTR was immunoprecipitated from detergent lysates with MrPink antibody and analyzed on 7.5% SDS-PAA gels (top panel). Remaining lysates were digested with 25 μg/mL ProtK for 15 min. Proteolytic fragments were immunoprecipitated with E1-22 antibody against TMD1, with 3G11 and 7D12 against different epitopes on NBD1, and with G449, which mainly detects RE and N-terminal residues of R (bottom panel) and analyzed by 12% SDS-PAGE. **D)** HEK293T expressing wild-type (wt) CFTR, F508del-CFTR, or F508del -CFTR with the R1102A mutation were radiolabeled for 15 min and chase for 2h (bottom panel) or not (0 h, top panel) in the presence of X307810, VX-445, or DMSO as control. CFTR was immunoprecipitated with MrPink antibody and analyzed by 7.5% SDS-PAGE.

We therefore transfected cells with CFTR expression constructs ΔN2, ΔN3, and ΔN4 missing the corresponding number of amino acids from the N-terminus. These cells were radiolabeled for 15 min and chased for 2 h in the presence of X307810, lysed and analyzed in the limited-proteolysis assay (Figure 5B). The previously defined T1d and T1e proteolytic fragments had the same mobility in the truncation mutants as in wild-type CFTR, because their N-terminal borders are at aa positions 36 and 17, respectively, C-terminal to the three truncations. In contrast, like T1f, T1g progressively became shorter and more sensitive to ProtK in mutants from which 2, 3, or 4 N-terminal residues were removed (Figure 5B, bottom gel panel and lane traces). Since the N-terminal border of T1f in wild-type CFTR is at the start methionine, we conclude that T1g has the same N-terminus as the T1f fragment.

We next determined the C-terminus of the T1g fragment in cells expressing wild-type CFTR. Cells were radiolabeled for 15 min and chased for 2 h in the presence of X307810, lysed and analyzed in the limited proteolysis assay (Figure 5C). Since T1g has the same N-terminus as T1f, yet is larger than T1f, and is detected by the MrPink antibody against NBD1, we inferred that it contains an epitope on NBD1. We immunoprecipitated proteolytic fragments from the lysates with antibodies that recognize different regions of NBD1, including monoclonal 3G11 (aa 396-405), monoclonal 7D12 (aa 531-540) and polyclonal G449 (aa 653-716) (Figure 5A). Only the monoclonal 3G11 detected the T1g fragment, while the antibodies against more C-terminal parts of NBD1 did not immunoprecipitate T1g. An estimation of the T1g molecular weight from multiple gels (Figs 2-5) uncovered that its C-terminus is located after the 3G11 epitope (aa 396-405) consistent with our previous finding that ProtK cleaves the N terminus of NBD1 in the intrinsically disordered Regulatory Insertion (aa 404-436) at position 428 (22). We concluded that X307810 extends the T1f fragment with C-terminal sequence, by protecting the linker between TMD1 and NBD1 up to the Regulatory Insertion.

## DISCUSSION

Pharmacological approaches for treatment of CF are centered on small-molecule modulators that improve folding of missense CFTR variants to an extent that these are licensed for export to the Golgi complex and rescued from degradation at the ER. This increases the number of channels on the cell surface. A critical pillar in this strategy evidently is corrector efficacy (36, 37).

Combinations of two corrector compounds in Trikafta, with distinct binding sites on CFTR and independent and additive effects on CFTR biogenesis, proved to be key to the initial success for CF patients carrying at least one F508del allele. The subsequent label expansion to other variants made a much more diverse group of patients eligible for the drug. Despite this major progress in CF care, the Trikafta correctors do not completely rescue biogenesis and fail to improve thermodynamic stability of F508del-CFTR (36, 38). Also, ∼10% of CF patients with missense mutations do not benefit from the presently available clinical compounds, and time will tell how well long-term treatment will be tolerated. This calls for development and characterization of additional and novel correctors with improved mode of action. Understanding how correctors work will also help dissect the molecular mechanisms underlying the folding of individual domains in CFTR and their assembly into a mature functional chloride channel.

The new corrector compounds we describe in this paper are built on the X339688 (compound 1) scaffold in which we have explored the chemical space of the cyclopropyl ring adjacent to the carbonyl group of the acylsulfonamide moiety. The most effective ones were X307810 and X323022, with EC_50_ values in the single digit nanomolar range as measured by TECC assays in HBE F508del/F508del cells. The activity of X307810 (S,R) was stereospecific, because its enantiomer, X339690 (S,S) had an 11-fold higher EC_50_ for rescue of F508del-CFTR function. Since X323022 and X307810 had very similar properties in rescue of F508del-CFTR from the ER and in functional assays, we focused on the characterization of X307810 as representative compound. Comparison of the effects of X307810 and the C1 corrector X281602 on rescue of F508del-CFTR from the ER showed that the high efficacy of X307810 cannot be attributed to partial C1 corrector behavior: the two compounds have additive modes of action on CFTR folding, and X307810 does not provoke the typical cotranslational biochemical response on TMD1 (T1aa) we have shown for C1 correctors X281602 (this manuscript), VX-661, and VX-809 (35). T1aa arises from protection from proteolysis around residue 50, caused by improved packing of the cytoplasmic N-terminus of TMD1 with ICL1 and the TMD1 C-terminus.

De-novo protein folding follows a vectorial pathway along meandering transient ‘detours’, including non-native intramolecular interactions, to a final folded state (39). Drug-induced conformational changes during early cotranslational stages of CFTR folding therefore invariably translate into downstream effects on domain assembly. Indeed, F508del-CFTR treated with the C1 corrector VX-661 is rescued indirectly by enhanced domain assembly (evidenced by limited-proteolysis fragments T1d-f, T2c, and N2a), due to improved cotranslational stability of TMD1 (T1aa) without rescue of the primary defect in NBD1 (Figure 4B). The T1d-f, T2c and N2a fragments derived from TMD1, TMD2, and NBD2, respectively, can only be generated from CFTR molecules that have completed cotranslational folding and have progressed through the posttranslational domain assembly stage. Class-2 correctors such as VX-445 enhance domain assembly directly, working largely posttranslationally. The limited-proteolysis assay gave the same phenotype for X307810-corrected F508del-CFTR in terms of the T1d-f, T2c, and N2a fragments. The new corrector compounds share an acylsulfonamide group with VX-445, which interacts with residues in the proximal part of the TMD1 lasso (S18, R21) and TM11 (W1098, R1102) of TMD2. We anticipated that the interaction of X307810 will depend in part on these amino acids as well. In accord, rescue of F508del-CFTR by X307810 indeed was prevented by mutation of R1102 (Figure 5D).

What makes X307810 distinct from other correctors including VX-445 is its unexpected protective effect on the linker between TMD1 and NBD1, extending into the N-terminal part of the regulatory insertion (RI). The TMD1-NBD1 linker becomes protected from ProtK in both F508del and wild-type CFTR when cells are treated with X307810. This results in T1g, a stable 40-kDa proteolytic fragment containing TMD1 in its entirety, extended C-terminally into the unstructured RI. Linkers connecting the domains in multidomain proteins provide structural flexibility and enhance concentration locally for facilitating (40) or limiting (41) domain-domain interactions. Linkers oftentimes have less secondary structure, which results in more accessible cleavage sites. Indeed, also in folded CFTR, the boundaries of limited-proteolysis fragments are just outside domains and coincide with linker regions (22). Cleavage between domains occurs before cleavage within domains. Much effort went into understanding the role of the dephosphorylated state of the linker region R as autoinhibitory regulator of the interaction between NBD1 and NBD2. Similarly, the role of intracellular loops in relaying signaling from the NBDs to movement of the transmembrane helices forming the ion channel is understood in some detail. In contrast, little is known about the linkers connecting TMD1 with NBD1 and TMD2 with NBD2, neither for CFTR nor for most other ABC transporters.

The decreased proteolysis of the TMD1-NBD1 linker is caused by a protection of the linker by an interacting protein sequence, either from within CFTR (domain folding or domain assembly) or from another protein. The interactor leaves a so-called footprint. CFTR has been found to interact with at least 80 proteins at different stages of its life (42), but the proteolytic change effected by X323022 and X307810 involves a late stage of its folding and the TMD1-NBD1 linker. This leaves only one identified candidate protein: the catalytic domain of PKA (PKA-C) (43). PKA activates CFTR to open its channel by phosphorylation of the R region (44, 45) and was shown to bind the CFTR N-terminus and TMD1-NBD2 linker (43). Whether PKA is involved in CFTR folding remains to be determined. A likely cause for linker protection is a X323022/X307810-corrector-induced conformational change in CFTR that leads to tight packing of the linker to TMD1 or NBD1.

The linker already was shown to contribute to a novel conformation identified in the Govaert lab. They recently discovered a novel conformation of NBD1 distinct from previous descriptions of the canonical NBD1 structure and extending the linker between TMD1 and NBD1 (46). In this so-called alternative β-SS conformation, the N-terminal antiparallel ABC-β subdomain (aa389-493) of NBD1 underwent architectural changes compared to the canonical structure. The N-terminal S1 beta strand (aa386-424) of RI now has become unstructured and the C-terminus of RI gained more structure through a new beta strand between aa 425-436. This extends the linker between TMD1 and NBD1 from 376-386 to 376-425. The β-SS conformation and canonical conformation are in equilibrium, with NBD1 reversibly sampling each of them.

In this alternative conformation, the linker between TMD1 and NBD1 is continuous with RI, the most N-terminal part in NBD1, and its modifications determine fate of CFTR. RI contains 3 phosphorylation sites (Thr421, Ser 422 and Ser427) that are phosphorylated 10x more abundantly in wild-type than in F508del-CFTR and it contains a Lys420 that is ubiquitinated in F508del-CFTR but methylated in wild-type CFTR (43). The extent of phosphorylation on Thr421, Ser 422, and Ser427 correlates with transport from ER to Golgi and is strongly decreased in N1303K and F508del-CFTR mutants (47). Inhibition of RI phosphorylation leads to ubiquitination of Lys420 (and Lys442), which marks CFTR for degradation. This reciprocal mechanism is consistent with the finding that deleson of RI improves maturason of F508del-CFTR (48), allowing escape from surveillance for degradason early in the folding pathway. Rescue of F508del-CFTR by deleson of RI hence may be due to post-translasonal modificasons, which likely have a basis in a conformasonal stabilizason.

We found that the impact of the new correctors on CFTR structure resulted in an extension of TMD1 limited-proteolysis fragments at the C-terminus from aa 383 for T1def to aa 428 in T1g, just beyond the 3G11 epitope in the N-terminus of RI. Clearly in the absence of correctors this region has a conformation that does not permit ProtK access to the 9 cleavage sites present in it. Given the stabilizing effect of the new corrector compounds on F508del and the pivotal role of the TMD1-NBD1-linker-RI sequence in CFTR proteostasis, we speculate that X323022 and X307810 mode of action may be through shifting the equilibrium between the canonical and alternative states the Govaert lab found towards the more stable conformation, leading to enhanced domain assembly without correction of NBD1 folding.

X323022 and X307810 have an overlapping binding site in TMD2 with VX-445, as all three require an arginine residue in position 1102, but the novel correctors demonstrate a distinct mode of action, more tightly packing the linker between TMD1 and NBD1 upon domain assembly. All three do not stabilize TMD1 alone against degradation, but require TMD2. The appearance of fragment T1g from wild-type CFTR upon treatment with X323022 or X307810 demonstrates action on the domain-assembled form. F508del-CFTR however does not assemble its domains without correctors but does respond strongly to these correctors by assembling its domains. Similarly, C1 correctors enhance domain assembly by stabilizing TMD1. We conclude that F508del (and probably most if not all CFTR missense mutants) do probe the assembled state but in an equilibrium strongly skewed towards the non-assembled stated. Correctors than change this equilibrium through various modes of action, and rescue the mutant protein, albeit not always to functional rescue. Defective domains are not corrected but the defect is compensated structurally in the assembled protein. This mechanism explains why by now 179 CF-disease-causing missense mutations have been FDA-approved for Trikafta. Domain assembly turned out to have been the holy grail of rescue, not domain correction per se, and X323022 or X307810 have shown that a single corrector is sufficient to rescue a large set of CFTR mutants.

## MATERIALS AND METHODS

### Synthesis and characterization of corrector compounds

X281602 and X281605 (C1-type correctors) have been described before as compound 10c and 10 respectively (29). Potentiator compounds X283614 and X283649 have been described as compound 4 and 5, respectively (29), and X289990 (C2-type corrector) is previously described compound 19 (29).

The synthesis and characterization of compound **1** is described in US patent 2019/0077784. The synthetic protocols for compounds **4** and **5** are detailed below and in Figure S1 (US patent 2022/0213041). The synthesis of compounds **2**, **3** and **6**–**8** was achieved in analogous fashion.

Compound **10**: *rac*-(1*r*,2*s*)-methyl 1-(2-methoxy-5-methylphenyl)-2-phenylcyclopropanecarboxylate. A solution of diazo methyl ester **9** (200 mg, 0.908 mmol) in dichloromethane (3 mL) was added over 4 hours by syringe pump to a solution of styrene (315 µL, 2.72 mmol, Aldrich) and rhodium(II) acetate dimer (2.0 mg, 4.5 µmol, Aldrich) in dichloromethane (6 mL) at ambient temperature. After stirring for an additional 12 hours, the reaction was concentrated under reduced pressure, and the crude residue was purified by flash chromatography (ISCO CombiFlash, 0-30% ethyl acetate / heptanes, 40 g RediSep^®^ gold silica column) to afford **10** (257 mg, 0.867 mmol, 95% yield). ^1^H NMR (500 MHz, CDCl_3_) *δ* ppm 7.06 – 6.97 (m, 3H), 6.97–6.89 (m, 2H), 6.82 – 6.71 (m, 2H), 6.42 (d, *J* = 8.2 Hz, 1H), 3.65 (s, 3H), 3.30 (s, 3H), 3.20 (dd, *J* = 9.3, 7.4 Hz, 1H), 2.23 (d, *J* = 0.8 Hz, 3H), 1.96 (dd, *J* = 9.3, 5.0 Hz, 1H), 1.82 (dd, *J* = 7.4, 5.0 Hz, 1H). MS(APCI+) *m/z* 297.4 (M+H)^+^.

Compound **11**: *rac*-(1*r*,2*s*)-1-(2-methoxy-5-methylphenyl)-2-phenylcyclopropanecarboxylic acid. Lithium hydroxide (205 mg, 8.57 mmol) was added to a solution of **10** (254 mg, 0.857 mmol) in dioxane (4.6 mL) and water (1.1 mL). The reaction mixture was then heated to 80 °C for 16 hours before being acidified with 1 M hydrochloric acid and extracted with ethyl acetate. The organic phase was washed with 1 M hydrochloric acid, brine, dried with MgSO_4_, filtered, and concentrated under reduced pressure to afford **11** (218 mg, 0.772 mmol, 90% yield). ^1^H NMR (400 MHz, dimethyl sulfoxide-*d*_6_) *δ* ppm 12.08 (s, 1H), 7.05 – 6.90 (m, 4H), 6.87 (dd, *J* = 8.6, 2.2 Hz, 1H), 6.83 – 6.76 (m, 2H), 6.49 (d, *J* = 8.2 Hz, 1H), 3.27 (s, 3H), 3.03 (dd, *J* = 9.1, 7.2 Hz, 1H), 2.16 (s, 3H), 1.87 (dd, *J* = 7.2, 4.9 Hz, 1H), 1.74 (dd, *J* = 9.2, 4.8 Hz, 1H). MS(APCI+) *m/z* 283.4 (M+H)^+^.

*Compounds **4** and **5**:* (1*S*,2*R*)-1-(2-methoxy-5-methylphenyl)-*N*-(2-methylquinoline-5-sulfonyl)-2-phenylcyclopropane-1-carboxamide and (1*R*,2*S*)-1-(2-methoxy-5-methylphenyl)-*N*-(2- methylquinoline-5-sulfonyl)-2-phenylcyclopropane-1-carboxamide. A mixture of compound **11** (100 mg, 0.354 mmol), 2-methylquinoline-5-sulfonamide (91 mg, 0.407 mmol, prepared as in WO2018154519 A1), 1-(3-dimethylaminopropyl)-3-ethylcarbodiimide hydrochloride (136 mg, 0.708 mmol), and 4-dimethylaminopyridine (56.3 mg, 0.460 mmol) in dichloromethane (3.5 mL) was stirred at ambient temperature. After 4 hours, the reaction was acidified with trifluoroacetic acid (136 µL, 1.77 mmol) and concentrated under reduced pressure. The crude residue was then purified by reverse-phase HPLC (Waters Xbridge Prep C18 column, 42 mL / minute, 5-95% acetonitrile / 0.1% trifluoroacetic acid in water) to afford the racemic mixture of the title compound (137 mg, 0.282 mmol, 79% yield). The racemic mixture (100 mg, 0.206 mmol) was then separated by preparative chiral supercritical fluid chromatography (ChiralPak IC column, 45% methanol / CO_2_, 80 g/minute). The fractions containing the first eluting peak were concentrated under reduced pressure to afford **5** (45.1 mg, 0.093 mmol, 45% yield). ^1^H NMR (400 MHz, dimethyl sulfoxide-*d*_6_) *δ* ppm 11.45 (s, 1H), 8.85 (d, *J* = 8.9 Hz, 1H), 8.34 – 8.15 (m, 2H), 7.89 (dd, *J* = 8.5, 7.4 Hz, 1H), 7.57 (d, *J* = 8.9 Hz, 1H), 7.03 – 6.84 (m, 5H), 6.78 – 6.67 (m, 2H), 6.44 (d, *J* = 8.3 Hz, 1H), 3.10 – 2.97 (m, 4H), 2.71 (s, 3H), 2.17 (s, 3H), 1.95 (s, 1H), 1.37 (dd, *J* = 9.2, 5.3 Hz, 1H). MS(APCI+) *m/z* 487.2 (M+H)^+^. The fractions containing the second eluting peak were concentrated under reduced pressure to afford **4** (45.1 mg, 0.093 mmol, 45% yield). ^1^H NMR (500 MHz, dimethyl sulfoxide-*d*_6_) *δ* ppm 11.48 (s, 1H), 8.85 (d, *J* = 8.9 Hz, 1H), 8.29 – 8.17 (m, 2H), 7.89 (dd, *J* = 8.5, 7.4 Hz, 1H), 7.57 (d, *J* = 8.9 Hz, 1H), 7.01 – 6.83 (m, 5H), 6.76 – 6.66 (m, 2H), 6.44 (d, *J* = 8.2 Hz, 1H), 3.17 (s, 2H), 3.08 – 2.96 (m, 4H), 2.72 (s, 3H), 2.17 (s, 3H), 1.95 (s, 1H), 1.37 (dd, *J* = 9.2, 5.3 Hz, 1H). MS(APCI+) *m/z* 487.2 (M+H)^+^.

Compound **2**: (1S,2S)-1-(2-methoxy-5-methylphenyl)-2-(methoxymethyl)-N-((2-methylquinolin-5- yl)sulfonyl)cyclopropane-1-carboxamide. ^1^H NMR (500 MHz, Chloroform-d) δ ppm 9.60 (s, 1H), 9.33 (d, *J* = 8.9 Hz, 1H), 8.71 (d, J = 8.5 Hz, 1H), 8.49 (dd, *J* = 7.5, 1.0 Hz, 1H), 8.20 (s, 1H), 8.00 (dd, J = 8.5, 7.5 Hz, 1H), 7.64 (d, *J* = 9.0 Hz, 1H), 7.20 (ddd, *J* = 8.3, 2.3, 0.9 Hz, 1H), 7.03 (s, 1H), 6.83 (d, *J* = 8.3 Hz, 1H), 3.62 (s, 3H), 3.12 (s, 4H), 3.01 (s, 3H), 2.65 (s, 1H), 2.31 (s, 3H), 2.13 (p, *J* = 7.1 Hz, 1H), 1.56 (dd, *J* = 9.2, 4.6 Hz, 1H); LCMS(APCI+) *m/z* 455.2 (M+H)^+^.

Compound **3:** (1R,2R)-1-(2-methoxy-5-methylphenyl)-2-(methoxymethyl)-N-((2-methylquinolin-5- yl)sulfonyl)cyclopropane-1-carboxamide. ^1^H NMR (400 MHz, DMSO-d6) δ ppm 11.43 (s, 1H), 8.91 (d, *J* = 8.9 Hz, 1H), 8.26 (ddd, *J* = 7.7, 3.3, 1.1 Hz, 2H), 7.93 (t, *J* = 8.0 Hz, 1H), 7.65 (d, *J* = 8.9 Hz, 1H), 7.11 (dd, *J* = 8.3, 2.2 Hz, 1H), 6.97 (d, *J* = 2.2 Hz, 1H), 6.79 (d, *J* = 8.3 Hz, 1H), 3.34 (s, 3H), 2.98 (s, 3H), 2.94 (dd, *J* = 10.5, 5.9 Hz, 1H), 2.75 (s, 3H), 2.43 (dd, *J* = 10.5, 7.8 Hz, 1H), 2.26 (s, 3H), 2.00 (tt, *J* = 8.9, 6.4 Hz, 1H), 1.27 (dd, *J* = 6.7, 5.0 Hz, 1H), 1.06 (dd, *J* = 9.1, 4.9 Hz, 1H); LCMS(APCI+) *m/z* 455.2 (M+H)^+^.

Compound **6**: (1S,2S)-1-(2-methoxy-5-methylphenyl)-N-((2-methylquinolin-5-yl)sulfonyl)-2-(pyridin- 2-yl)cyclopropane-1-carboxamide. ^1^H NMR (400 MHz, dimethyl sulfoxide-*d*_6_) δ ppm 8.85 (d, *J* = 8.8 Hz, 1H), 8.17 (dd, *J* = 7.8, 3.3 Hz, 2H), 8.09 (dd, *J* = 5.2, 1.8 Hz, 1H), 7.84 (dd, *J* = 8.5, 7.4 Hz, 1H), 7.53 (d, *J* = 8.9 Hz, 1H), 7.43 (td, *J* = 7.7, 1.8 Hz, 1H), 6.96 (ddd, *J* = 7.6, 4.9, 1.1 Hz, 1H), 6.91 – 6.78 (m, 3H), 6.40 (d, *J* = 8.3 Hz, 1H), 3.17 (dd, *J* = 8.8, 6.9 Hz, 1H), 3.04 (s, 3H), 2.70 (s, 3H), 2.16 (s, 4H), 1.47 (dd, *J* = 8.9, 4.4 Hz, 1H). MS(APCI+) *m/z* 488.4 (M+H)^+^.

Compound **7**: (1S,2S)-1-(2-methoxy-5-methylphenyl)-2-(6-methylpyridin-2-yl)-N-((2-methylquinolin- 5-yl)sulfonyl)cyclopropane-1-carboxamide. ^1^H NMR (500 MHz, CDCl_3_) δ ppm 9.75 (d, *J* = 8.9 Hz, 1H), 8.75 (d, *J* = 8.4 Hz, 1H), 8.60 (d, *J* = 7.4 Hz, 1H), 8.05 (t, *J* = 7.9 Hz, 1H), 7.79 (d, *J* = 8.9 Hz, 1H), 7.70 (t, *J* = 7.9 Hz, 1H), 7.29 (d, *J* = 4.0 Hz, 1H), 7.14 (d, *J* = 2.2 Hz, 1H), 7.05 (dd, *J* = 8.5, 2.1 Hz, 1H), 6.43 (dd, *J* = 8.2, 4.4 Hz, 2H), 3.80 (t, *J* = 7.8 Hz, 1H), 3.29 (s, 3H), 3.07 (s, 3H), 2.75 (s, 3H), 2.28 (s, 3H), 2.17 2.04 (m, 2H). MS(APCI+) *m*/*z* 502.5 (M+H)^+^.

Compound **8**: (1S,2S)-2-(6-ethoxypyridin-2-yl)-1-(2-methoxy-5-methylphenyl)-N-((2-methylquinolin- 5-yl)sulfonyl)cyclopropane-1-carboxamide. ^1^H NMR (600 MHz, dimethyl sulfoxide-*d*_6_) δ ppm 11.40 (s, 1H), 8.83 (d, *J* = 8.8 Hz, 1H), 8.25 (d, *J* = 7.8 Hz, 2H), 7.90 (dd, *J* = 8.3, 7.5 Hz, 1H), 7.58 (d, *J* = 8.9 Hz, 1H), 7.33 (dd, *J* = 8.1, 7.3 Hz, 1H), 7.00 (s, 1H), 6.96 – 6.87 (m, 1H), 6.69 (d, *J* = 7.3 Hz, 1H), 6.44 (d, *J* = 8.3 Hz, 1H), 6.25 (dd, *J* = 8.2, 0.7 Hz, 1H), 3.84 (dq, *J* = 10.6, 7.1 Hz, 1H), 3.59 (dq, *J* = 10.7, 7.1 Hz, 1H), 3.07 (dd, *J* = 8.8, 6.9 Hz, 1H), 3.02 (s, 3H), 2.71 (s, 3H), 2.19 (s, 4H), 1.43 (dd, *J* = 8.8, 4.3 Hz, 1H), 1.08 (t, *J* = 7.1 Hz, 3H). MS(APCI+) *m/z* 532.4 (M+H)^+^.

### Cell lines and transfection

HEK293T cells (ATCC; CRL-3216) were cultured in DMEM, 10% FBS, 2 mM Glutamax-I (Life Technologies) at 37°C and 5% CO_2_. Cells were grown to 60-70% confluency and transfected in 6-cm dishes using polyethylenimine (PEI; Polysciences) as described (49). After 4 h, medium was exchanged, and cells were grown for 24 h before being used in experiments. CFBE41o- cells stably expressing F508del-CFTR were a gift from Dr. Bob Bridges (Rosalind Franklin University of Medical Sciences, North Chicago, IL). Cells were maintained in MEM (Gibco) supplemented with 10% FBS (Gibco), 1% penicillin/streptomycin (Gibco) and 500 µg/ml geneticin (Gibco) at 37°C with 5% CO_2_. Primary HBE cells from CFTR patients with homozygous F508del/F508del mutation (HBE F508del/F508del cells) used to generate table 1 transepithelial current clamp (TECC) data were obtained from Dr. Bob Bridges (Rosalind Franklin University of Medical Sciences, North Chicago, IL) (Donor ID BrCF001), the Marsico Lung Institute Tissue Procurement and Cell Culture Core at the University of North Carolina at Chapel Hill (Donor ID UNC48N), and McGill University (Donor ID McGill683). HBE F508del/F508del cells from Donor IDs BrCF001 and UNC25O were used in Figures 1C and 1D. Cells were seeded onto 24-well transwell inserts (Corning) coated with conditioned medium from cultured 3T3 fibroblasts and grown at an air liquid interface for 35 days in differentiation medium containing 2% Ultroser-G.

### Antibodies and cDNAs

The polyclonal rabbit antibodies E1-22 and TMD1-C against TMD1, MrPink against NBD1, TMD2-C against TMD2, rabbit antibody G449 against RE-R (thanks to Dr. Angus Nairn (Yale University, New Haven CT USA)), mouse monoclonal antibody 596 against NBD2 (thanks to Dr. John Riordan (University of North Carolina, Chapel Hill NC USA), and the Cystic Fibrosis Foundation (CFF)), 7D12 (thanks to Dr. Philip J. Thomas, (University of Texas Southwestern Medical Center, Dallas TX USA), and 3G11 (thanks to Dr. William Balch (Scripps Research, La Jolla CA USA)) against NBD1, and anti-actin antibody have been described (22). GFP antibodies were generated against recombinant GST- BFP (50). The monoclonal antibody against Na^+^/K^+^ ATPase antibody was from Abcam and IRDye 800CW labeled secondary antibody goat anti-mouse was from LI-COR. CFTR expression constructs in pBi-CMV2 were described before (22, 35, 49, 51).

### Cell-surface expression-Horse Radish Peroxidase (CSE-HRP) assay

To assay the effect of correctors on F508del-CFTR transport to the plasma membrane, the CSE-HRP assay was performed as described (52). Briefly, CFBE41o- cells stably expressing F508del-CFTR along with HRP were plated in 384-well plates and incubated at 37°C with 5% CO_2_ for 72 hours in the presence of 0.5 µg/mL doxycycline to induce F508del-CFTR-HRP expression. Cells were then incubated with corrector compounds for 18-24 hours at 33 °C. Following incubation, plates were washed with Dulbecco’s phosphate buffered saline (DPBS) and then 50 µL of the HRP substrate, Luminol, were added. HRP activity was measured using EnVision® Multilabel Plate Reader (Perkin Elmer).

### Pulse-chase assay

De-novo protein synthesis was studied using pulse-chase analysis, as described (22, 51). Briefly, cells were starved for 15 minutes of methionine and cysteine (DMEM depletion medium; Gibco Life Technologies) and then pulse labeled for 15 minutes with 132 µCi/60 mm dish of EasyTag Express ^35^S protein labeling mix (Perkin Elmer) in DMEM depletion medium. After radiolabeling, medium was exchanged, and cells were incubated in growth medium containing 5 mM methionine and cysteine for the indicated chase times. Cells were lysed on ice in 1% Triton X-100, 20 mM MES, 100 mM NaCl, 30 mM Tris/HCl pH 7.4 (MNT-Triton), followed by limited proteolysis and/or immunoprecipitation of CFTR by the MrPink antibody. Compounds were added during starvation, pulse labeling, chase and cell lysis.

### Limited proteolysis, immunoprecipitation, and analysis on SDS-PAGE

After the pulse chase, cells were lysed and limited proteolysis was done as described before (22). Briefly, detergent cell lysate was incubated with 25 µg/ml Proteinase K (ProtK) (Sigma-Aldrich) for 15 minutes on ice to generate protein fragments. The reaction was stopped by addition of lysis buffer containing 2 mM PMSF and 5 µg/ml of chymostatin, leupeptin, antipain, pepstatin (CLAP, all purchased from Sigma-Aldrich). Lysates then were subjected to immunoprecipitation using CFTR domain-specific antibodies: E1-22 for TMD1 fragments, MrPink for NBD1 fragments, TMD2-C for TMD2 and 596 for NBD2 as described (22). Proteins were analyzed by SDS-PAGE: 7.5% for full-length CFTR, 12% for proteolyzed fragments of CFTR. Gels were dried and exposed to phosphor screens for autoradiography (Typhoon FLA 7000 GE Healthcare).

### Western-blot analysis of F508del-CFTR expression

CFBE41o- cells stably expressing F508del-CFTR were seeded overnight at 0.75×10^6^ cells/well onto six-well dishes and then treated with compounds for 24 h. Cells were lysed in RIPA buffer (Sigma) containing protease inhibitor cocktail (Roche). Equal amounts of cleared lysate protein were loaded onto (3-8%) NuPAGE Tris-acetate gels (Invitrogen) and separated for 2 h 20 min at 150 V. Proteins were transferred onto nitrocellulose membranes using Novex Semi-Dry Blotter (Invitrogen). Membranes were probed overnight with mouse anti CFTR 596 antibody (1:5,000) and mouse anti- Na^+^/K^+^ ATPase (1:5,000). Membranes were then probed for 1 h with IRDye 800CW goat anti-mouse IgG (1:15,000), imaged on a LI-COR Odyssey system. Western blots of lysates from transfected HEK293T cells transiently expressing F508del-CFTR and treated for 24 h with corrector compounds were done as described (49).

### Transepithelial current clamp (TECC) assay

Mucus was removed from the apical side of HBE F508del/F508del cells 72 h prior to functional measurements by incubating cells at the apical side for 30 min with 3 mM DTT in Dulbecco’s PBS with Ca^2+^ and Mg^2+^. Mucus was aspirated and cells were washed once more and then treated with compounds at the basolateral side for 18-24 at 37°C with 5% CO_2_. CFTR activity was measured as CFTR equivalent current (I_eq_) generated by HBE cells in 24 channel electrode TECC assays. Prior to TECC measurement, cells were transferred into a bicarbonate- and serum-free F12 medium containing the desired compounds. The media was added to both apical and basolateral sides of the transwell inserts and plates then were equilibrated at 37°C for 30 min in a CO_2_-free incubator. Following equilibration, 8 initial measurements at ∼2-min intervals were recorded to determine the baseline resistance. Additionally, the transepithelial resistance was measured after the sequential addition of a) 6 µM benzamil at the apical side to inhibit the ENaC channel, b) 10 µM of forskolin at the apical and basolateral sides to activate the CFTR channel, and c) 20 µM bumetanide at the basolateral side to inhibit the Na:2Cl:K cotransporter and 20 µM CFTRinh-172/GlyH-101 at the apical side to inhibit the CFTR channel.

### Data analysis

Results from CSE-HRP assays were analyzed using Accelrys® Assay Explorer v3.3 and EC_50_ and the maximum-%-activity were calculated as described (52, 53). Transepithelial conductance (G_t_) was calculated from resistance measurements corrected for the combined solution series and empty insert resistances (7, 53). G_t_ values along with the transepithelial potential difference (V_t_) values corrected for the electrode offset potential were used to calculate I_eq_ using Ohm’s law (I_eq_ = V_t_. G_t_). The area under the curve (AUC) for the period between the forskolin peak-I_eq_ response and inhibitor addition was calculated using a one-third trapezoid method. I_eq_ and AUC values were fitted to a sigmoidal curve equation with Hill coefficient 1 using GraphPad Prism V9.5. Western blots were analyzed with proprietary Licor software and background subtraction was done with the box method. Quantitation and lane scans of radiolabeling experiments of full-length CFTR and domain fragments were done by phospohorimaging in ImageQuant 8.1 (GE-Healthcare) according to manufacturer specifications. Background subtraction on the raw values was done using the rolling disk using a disk size of 10,000 and was compared to the raw data to validate the quantitation and background subtraction. Protein-structure figures were prepared using ChimeraX (54), or Pymol. All graphs were made using Spotfire (Tibco, Perkin Elmer). Results are expressed as mean ± S.D. or S.E.M as indicated. Statistical analyses were carried out using GraphPad Prism V9.5.

## ACKNOWLEDGMENTS

This work was supported by The Cystic Fibrosis Foundation (BRAAKM08XX0, BRAAKM14XX0, BRAAKM18G0 to IB), grants from the Innovation fund for Chemistry (LIFT and TKI Holland Chemistry to PvdS and IB), the Netherlands Cystic Fibrosis Foundation (HIT-CF 3.0 to IB), Stichting Zeldzame Ziekten Fonds via Stichting Muco & Friends, the Netherlands Organization for Health Research and Development (Zon-MW Top 40-00812-98-14103), the UK Cystic Fibrosis Trust (to IB), and Gravity consortium grant 2024 to IB and PvdS. We are grateful to René Scriwanek and Dr. Ilias Balourdas for artwork, and we thank members of the Braakman-Van der Sluijs lab for discussions.

## REFERENCES

1. M. J. Welsh, A. E. Smith, Molecular mechanisms of CFTR chloride channel dysfunction in cystic fibrosis. Cell 73, 1251–1254 (1993).

2. A. Alam, K. P. Locher, Structure and Mechanism of Human ABC Transporters. Annu Rev Biophys 52, 275–300 (2023).

3. R. C. Ford, D. Marshall-Sabey, J. Schuetz, Linker Domains: Why ABC Transporters ‘Live in Fragments no Longer’. Trends Biochem Sci 45, 137–148 (2020).

4. W. R. Thelin et al., Direct interaction with filamins modulates the stability and plasma membrane expression of CFTR. J Clin Invest 117, 364–374 (2007).

5. S. H. Cheng et al., Phosphorylation of the R domain by cAMP-dependent protein kinase regulates the CFTR chloride channel. Cell 66, 1027–1036 (1991).

6. J. E. Ideozu et al., Diversity of CFTR variants across ancestries characterized using 454,727 UK biobank whole exome sequences. Genome Med 16, 43 (2024).

7. H. Bihler et al., In vitro modulator responsiveness of 655 CFTR variants found in people with cystic fibrosis. J Cyst Fibros 10.1016/j.jcf.2024.02.006 (2024).

8. H. Grasemann, F. Ratjen, Cystic Fibrosis. N Engl J Med 389, 1693–1707 (2023).

9. J. R. Riordan et al., Identification of the cystic fibrosis gene: cloning and characterization of complementary DNA. Science 245, 1066–1073 (1989).

10. P. van der Sluijs, H. Hoelen, A. Schmidt, I. Braakman, The folding pathway of ABC transporter CFTR: effective and robust. J Mol Biol 10.1016/j.jmb.2024.168591, 168591 (2024).

11. W. E. Balch, D. M. Roth, D. M. Hutt, Emergent properties of proteostasis in managing cystic fibrosis. Cold Spring Harb Perspect Biol 3 (2011).

12. T. Okiyoneda et al., Peripheral protein quality control removes unfolded CFTR from the plasma membrane. Science 329, 805–810 (2010).

13. P. R. Burgel, E. Burnet, L. Regard, C. Martin, The Changing Epidemiology of Cystic Fibrosis: The Implications for Adult Care. Chest 163, 89–99 (2023).

14. F. Van Goor et al., Rescue of CF airway epithelial cell function in vitro by a CFTR potentiator, VX-770. Proc Natl Acad Sci U S A 106, 18825–18830 (2009).

15. D. Keating et al., VX-445-Tezacaftor-Ivacaftor in Patients with Cystic Fibrosis and One or Two Phe508del Alleles. N Engl J Med 379, 1612–1620 (2018).

16. K. Fiedorczuk, J. Chen, Molecular structures reveal synergistic rescue of Δ508 CFTR by Trikafta modulators. Science 378, 284–290 (2022).

17. K. Fiedorczuk, J. Chen, Mechanism of CFTR correction by type I folding correctors. Cell 185, 158–168.e111 (2022).

18. O. Laselva et al., Rescue of multiple class II CFTR mutations by elexacaftor+tezacaftor+ivacaftor mediated in part by the dual activities of elexacaftor as both corrector and potentiator. Eur Respir J 57 (2021).

19. Y. Ivanenkov et al., The Hitchhiker’s Guide to Deep Learning Driven Generative Chemistry. ACS Med Chem Lett 14, 901–915 (2023).

20. A. Tropsha, O. Isayev, A. Varnek, G. Schneider, A. Cherkasov, Integrating QSAR modelling and deep learning in drug discovery: the emergence of deep QSAR. Nat Rev Drug Discov 23, 141–155 (2024).

21. F. Liu et al., Structure-based discovery of CFTR potentiators and inhibitors. Cell 187, 3712–3725.e3734 (2024).

22. J. Im et al., ABC-transporter CFTR folds with high fidelity through a modular, stepwise pathway. Cell Mol Life Sci 80, 33 (2023).

23. N. Soya et al., Folding correctors can restore CFTR posttranslational folding landscape by allosteric domain-domain coupling. Nat Commun 14, 6868 (2023).

24. E. Hong, A. Shi, P. Beringer, Drug-drug interactions involving CFTR modulators: a review of the evidence and clinical implications. Expert Opin Drug Metab Toxicol 19, 203–216 (2023).

25. D. Purkayastha et al., Drug-drug interactions with CFTR modulator therapy in cystic fibrosis: Focus on Trikafta®/Kaftrio®. J Cyst Fibros 22, 478–483 (2023).

26. C. J. Bathgate et al., Positive and negative impacts of elexacaftor/tezacaftor/ivacaftor: Healthcare providers’ observations across US centers. Pediatr Pulmonol 58, 2469–2477 (2023).

27. X. Wang et al., Discovery of 4-[(2R,4R)-4-({[1-(2,2-Difluoro-1,3-benzodioxol-5- yl)cyclopropyl]carbonyl}amino)-7-(difluoromethoxy)-3,4-dihydro-2H-chromen-2-yl]benzoic Acid (ABBV/GLPG-2222), a Potent Cystic Fibrosis Transmembrane Conductance Regulator (CFTR) Corrector for the Treatment of Cystic Fibrosis. J Med Chem 61, 1436–1449 (2018).

28. K. E. Oliver et al., Slowing ribosome velocity restores folding and function of mutant CFTR. J Clin Invest 129, 5236–5253 (2019).

29. X. Wang, C. Tse, A. Singh, Discovery and Development of CFTR Modulators for the Treatment of Cystic Fibrosis. J Med Chem 68, 2255–2300 (2025).

30. P. R. Sosnay et al., Defining the disease liability of variants in the cystic fibrosis transmembrane conductance regulator gene. Nat Genet 45, 1160–1167 (2013).

31. F. Van Goor, H. Yu, B. Burton, B. J. Hoffman, Effect of ivacaftor on CFTR forms with missense mutations associated with defects in protein processing or function. J Cyst Fibros 13, 29–36 (2014).

32. F. Anglès, C. Wang, W. E. Balch, Spatial covariance analysis reveals the residue-by-residue thermodynamic contribution of variation to the CFTR fold. Commun Biol 5, 356 (2022).

33. H. Y. Ren et al., VX-809 corrects folding defects in cystic fibrosis transmembrane conductance regulator protein through action on membrane-spanning domain 1. Mol Biol Cell 24, 3016–3024 (2013).

34. M. Pizzonero et al., Discovery of GLPG2737, a Potent Type 2 Corrector of CFTR for the Treatment of Cystic Fibrosis in Combination with a Potentiator and a Type 1 Co-corrector. J Med Chem 67, 5216–5232 (2024).

35. B. Kleizen et al., Co-Translational Folding of the First Transmembrane Domain of ABC-Transporter CFTR is Supported by Assembly with the First Cytosolic Domain. J Mol Biol 433, 166955 (2021).

36. G. Veit et al., Structure-guided combination therapy to potently improve the function of mutant CFTRs. Nat Med 24, 1732–1742 (2018).

37. V. Marchesin et al., A uniquely efficacious type of CFTR corrector with complementary mode of action. Sci Adv 10, eadk1814 (2024).

38. G. Veit, et al., Allosteric folding correction of F508del and rare CFTR mutants by elexacaftor- tezacaftor-ivacaftor (Trikafta) combination. JCI Insight 5 (2020).

39. A. Jansens, E. van Duijn, I. Braakman, Coordinated nonvectorial folding in a newly synthesized multidomain protein. Science 298, 2401–2403 (2002).

40. Q. Huang, M. Li, L. Lai, Z. Liu, Allostery of multidomain proteins with disordered linkers. Curr Opin Struct Biol 62, 175–182 (2020).

41. G. K. Schuurman-Wolters, M. de Boer, M. K. Pietrzyk, B. Poolman, Protein Linkers Provide Limits on the Domain Interactions in the ABC Importer GlnPQ and Determine the Rate of Transport. J Mol Biol 430, 1249–1262 (2018).

42. S. Pankow et al., ΔF508 CFTR interactome remodelling promotes rescue of cystic fibrosis. Nature 528, 510–516 (2015).

43. K. Fiedorczuk et al., The structures of protein kinase A in complex with CFTR: Mechanisms of phosphorylation and noncatalytic activation. Proc Natl Acad Sci U S A 121, e2409049121 (2024).

44. V. Kanelis, R. P. Hudson, P. H. Thibodeau, P. J. Thomas, J. D. Forman-Kay, NMR evidence for differential phosphorylation-dependent interactions in WT and DeltaF508 CFTR. Embo j 29, 263–277 (2010).

45. C. Marasini, L. Galeno, O. Moran, A SAXS-based ensemble model of the native and phosphorylated regulatory domain of the CFTR. Cell Mol Life Sci 70, 923–933 (2013).

46. D. Scholl et al., A topological switch in CFTR modulates channel activity and sensitivity to unfolding. Nat Chem Biol 17, 989–997 (2021).

47. S. Pankow, C. Bamberger, J. R. Yates, 3rd, A posttranslational modification code for CFTR maturation is altered in cystic fibrosis. Sci Signal 12 (2019).

48. A. A. Aleksandrov et al., Regulatory insertion removal restores maturation, stability and function of DeltaF508 CFTR. J Mol Biol 401, 194–210 (2010).

49. T. Hillenaar, J. Beekman, P. van der Sluijs, I. Braakman, Redefining Hypo- and Hyper-Responding Phenotypes of CFTR Mutants for Understanding and Therapy. Int J Mol Sci 23 (2022).

50. M. C. Hagemeijer et al., Membrane rearrangements mediated by coronavirus nonstructural proteins 3 and 4. Virology 458-459, 125-135 (2014).

51. M. van Willigen et al., Folding-function relationship of the most common cystic fibrosis-causing CFTR conductance mutants. Life Sci Alliance 2 (2019).

52. A. K. Singh et al., Biological Characterization of F508delCFTR Protein Processing by the CFTR Corrector ABBV-2222/GLPG2222. J Pharmacol Exp Ther 372, 107–118 (2020).

53. C. B. Vu et al., Synthesis and Characterization of Fatty Acid Conjugates of Niacin and Salicylic Acid. J Med Chem 59, 1217–1231 (2016).

54. E. C. Meng et al., UCSF ChimeraX: Tools for structure building and analysis. Protein Sci 32, e4792 (2023).

