## supplementary information for "A new corrector rescues F508del-CFTR folding through stabilization of the TMD1-NBD1 linker"

### SUPPLEMENTARY FIGURE LEGENDS

***Fig. S1: Synthesis of compounds 4 and 5***

Synthetic route for compounds **4** and **5**.

***Figure S2: The new compounds act additively with a C1 corrector***

**A)** HEK293T cells expressing CFTR508del-CFTR were treated overnight with 3 μM X281602 in combination with (or without) 3 μM of each X307810, X339689, X339690, and X323022 and analyzed by Western blot using antibodies against CFTR, actin, and GFP (transfection marker of pBi-cmv2 CFTRF508del plasmid). Positions of oligomannose ER form (ER) and complex-glycosylated Golgi form (G) are indicated. **B)** Bar graph with quantitation of % F508del-CFTR transported to the Golgi complex (G*100/(G+ER)). Data represented are means ± SD, of 3 experiments. Statistics analyze effects of C2 corrects in relation to DMSO and their effects on top of C1 corrector X281602. **C)** Bar graphs from panel B. Statistics analyze effects of C1 corrector X281602 on top of each C2 corrector, and comparison of C1 X281602 + C2 corrrectors X307810 or X323022 with the Vertex corrector combination in Trikafta, VX-445 + VX-661. *, **, ***, and **** indicate $p < 0.05$, $p < 0.005$, $p < 0.001$, $p < 0.0001$, respectively in one-tailed t tests).

***Fig. S3: X307810 does not have C1-corrector activity***

**A)** HEK293T cells expressing TMD1 were treated with corrector compound X307810 at 3 μM during starvation, radiolabeling, and chase. Cells were pulse labeled for 15 minutes and chased for up to 4 h in the presence of 10 μM cycloheximide. CFTR was immunoprecipitated using E1-22 and immunoprecipitates were resolved by 12.5% SDS-PAGE. **B)** HEK293T cells expressing wild-type (wt) CFTR or F508del-CFTR were treated with corrector compound X307810 at 3 μM final concentration during starvation and radiolabeling. Cells were pulse labeled for 15 minutes and lysed; CFTR was immunoprecipitated using E1-22 and immunoprecipitates were resolved by 7.5% SDS-PAGE (top panel). Remaining lysates were subjected to limited proteolysis for 15 min with 25 µg/mL ProtK. Protease-resistant fragments were immunoprecipitated with E1-22 and resolved by 12% SDS-PAGE (bottom panel). Band intensities of the T1a region were scanned to highlight the increase in TMD1-derived T1aa proteolytic fragment in the wild-type and F508del-CFTR proteolytic digests upon treatment with C1 corrector.

***Fig. S4: Milder proteolysis conditions uncover T1g in F508del-CFTR***

**A)** HEK293T cells expressing wild-type (wt) or F508del-CFTR were treated with modulator compound X307810 at 3 μM during starvation, radiolabeling, and chase. Cells were pulse labeled for 15 minutes and chased for 2 h or not (0 h). CFTR was immunoprecipitated using MrPink and immunoprecipitates were resolved by 7.5% SDS-PAGE. **B)** Remaining lysates were subjected to limited proteolysis for 15 min with a dilution series of ProtK ranging from 25 (0), 2.5 (1), 0.25 (2), 0.025 (3), to 0.0025 µg/mL (4) K. Protease-resistant fragments were immunoprecipitated with E1-22 and resolved by 12% SDS-PAGE. **C)** Same as be **B)** but limited proteolysis now was with the standard 25 µg/mL but for 1 (instead of 15) min.

***Fig. S5: NBD1-derived fragment gels for ProtK limited-proteolysis time course***

(**A, B**) HEK293T cells expressing wild-type (wt) or F508del-CFTR were labeled for 15 minutes and then chased for 2 h or not (0 h). Cells were incubated with corrector compound X307810 at 3 µM final concentration during starvation, radiolabeling, and chase. CFTR was immunoprecipitated from detergent lysates in parallel with E1-22 and MrPink antibodies and analyzed on 7.5% SDS-PAA gels. **(C, D)** Remainder of lysates was treated as in Figure 3D,E, except that samples were immunoprecipitated with NBD1 antibody.

Supplemental Figure-1

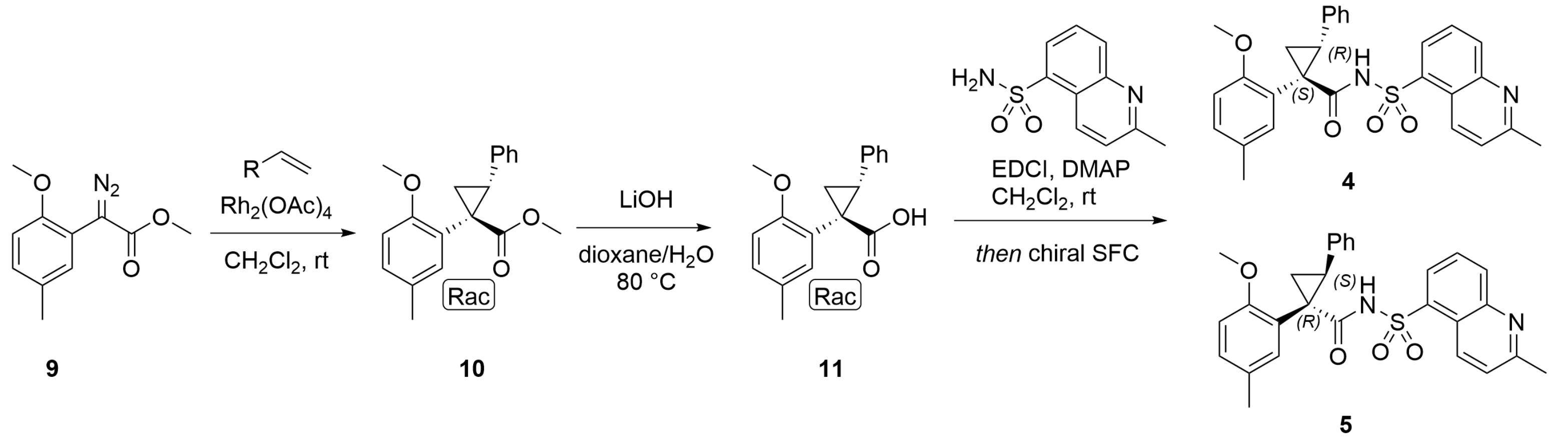

Supplemental Figure-2

(A)

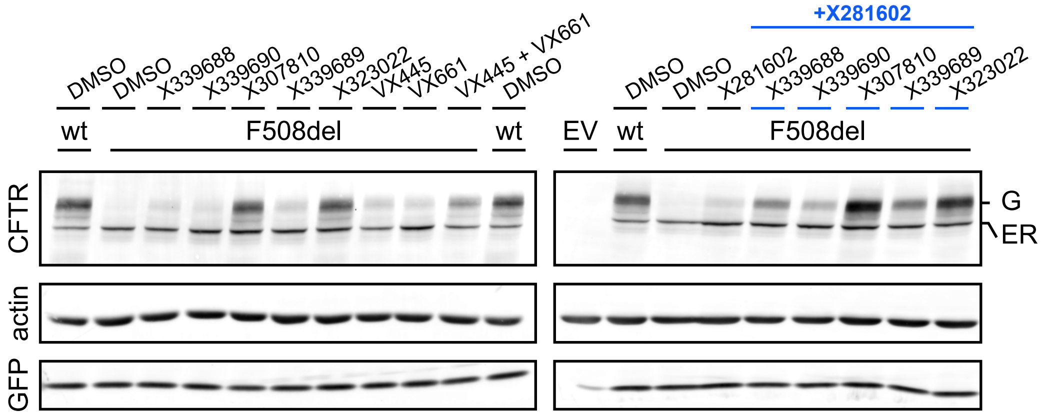


(B)

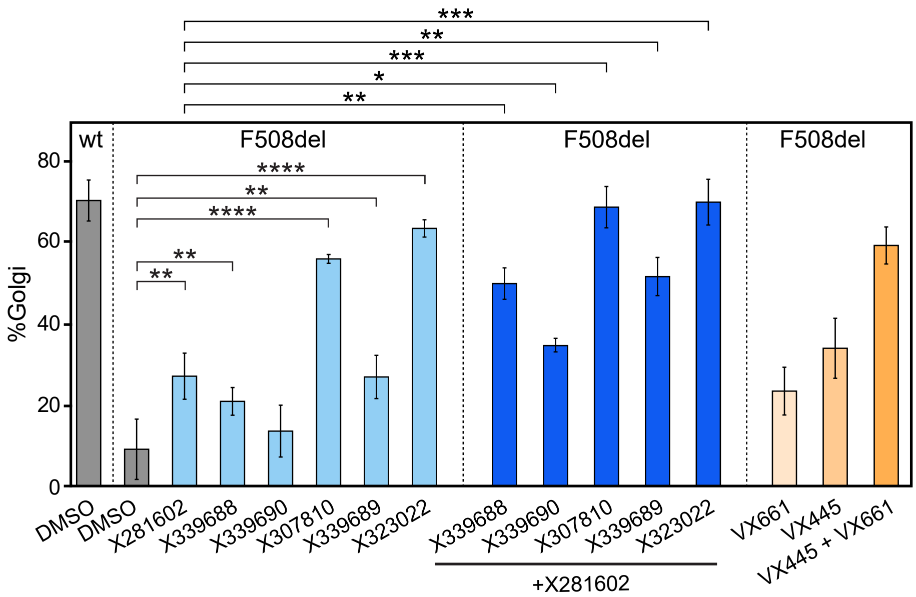


(C)

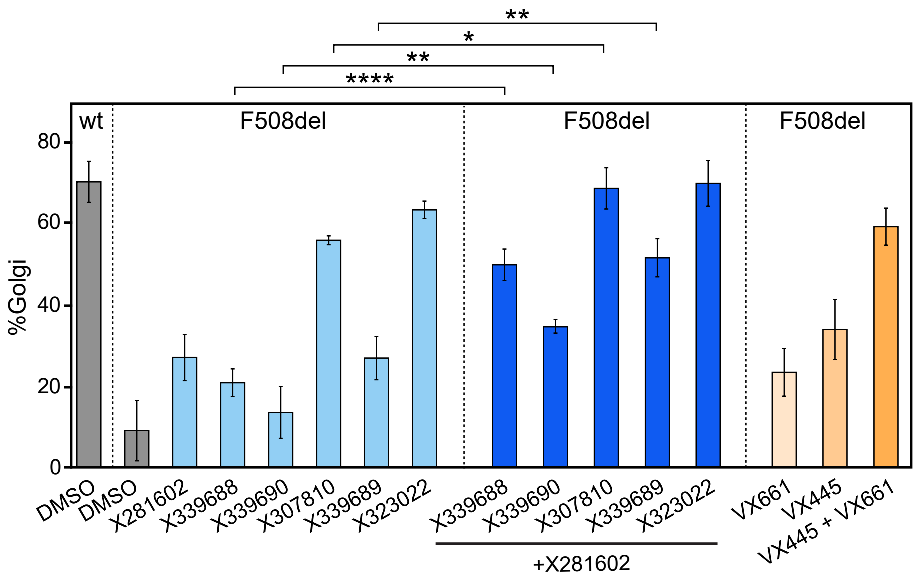

Supplemental Figure-3

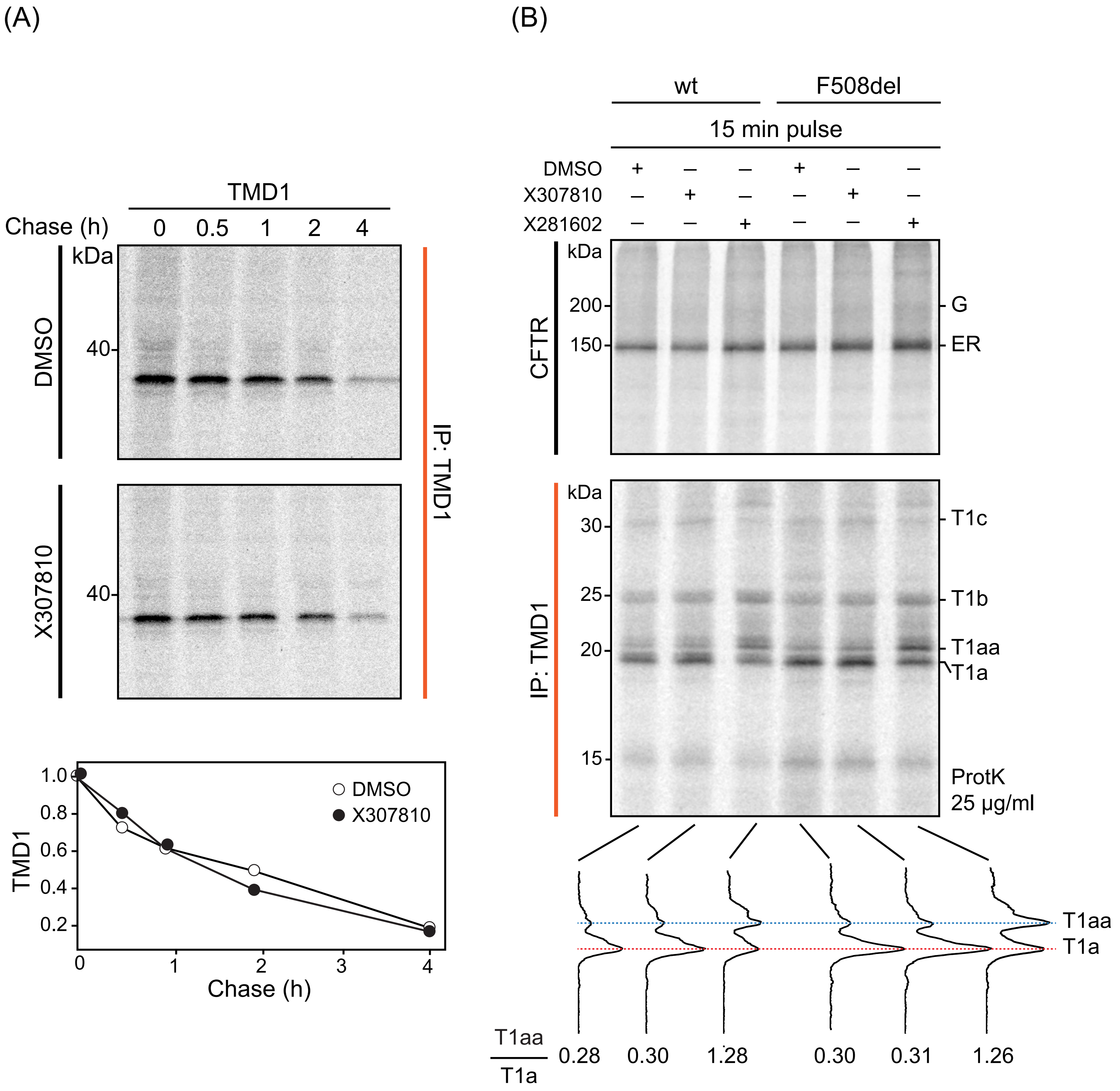

Supplemental Figure-4

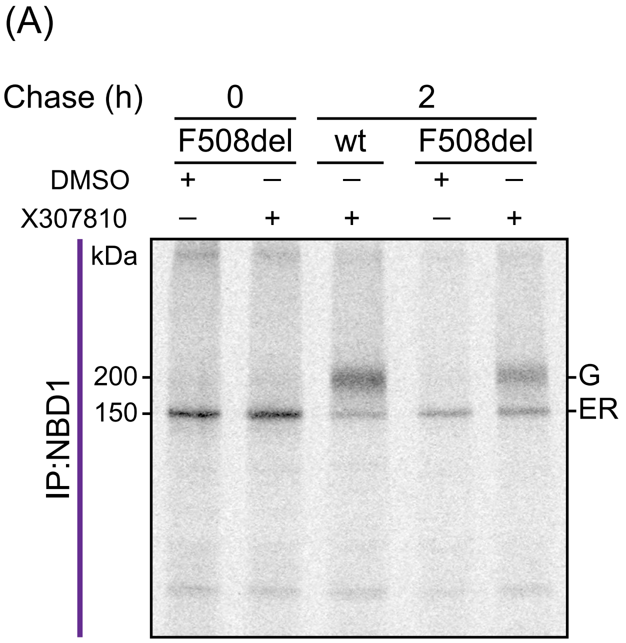


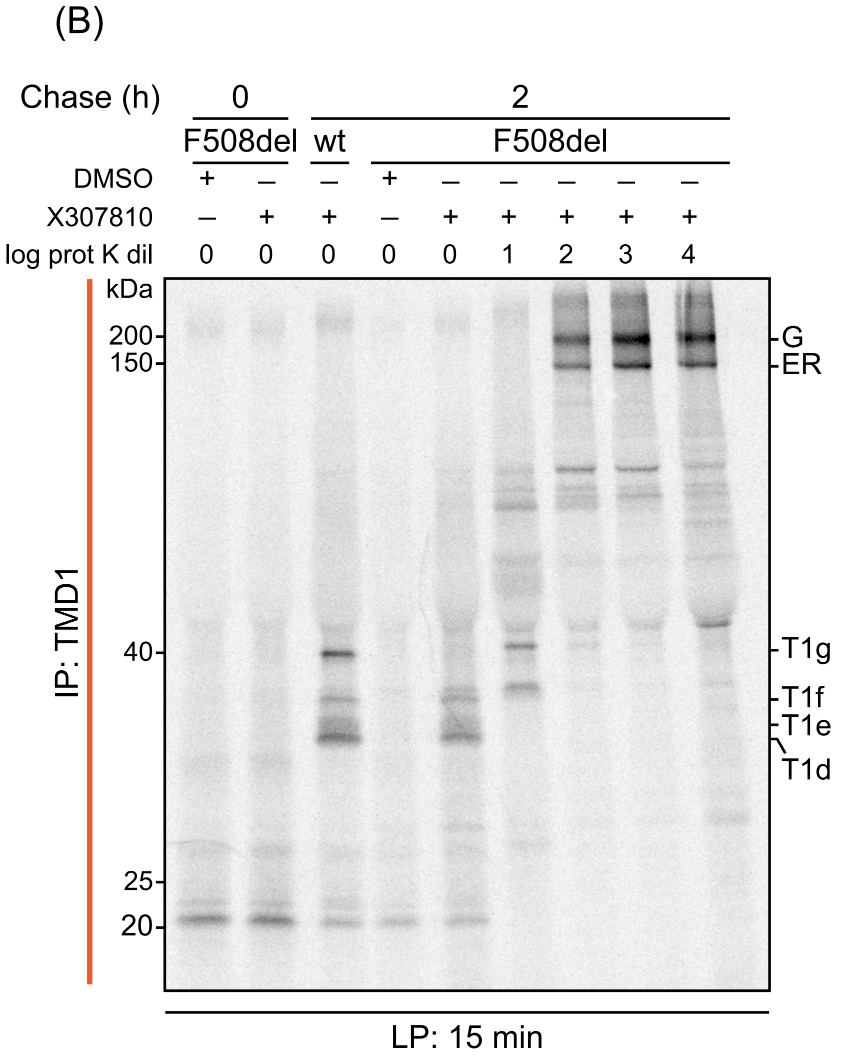


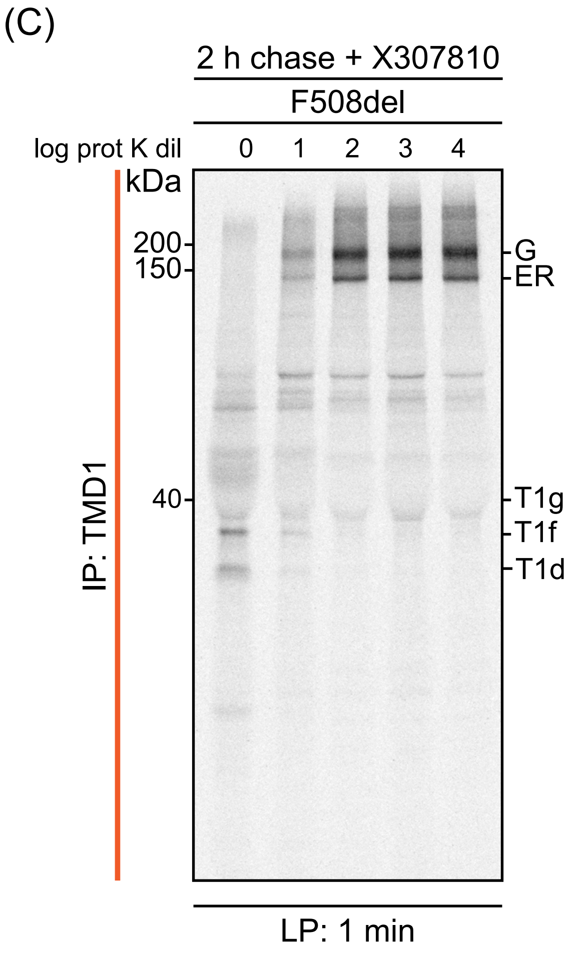

Supplemental Figure-5

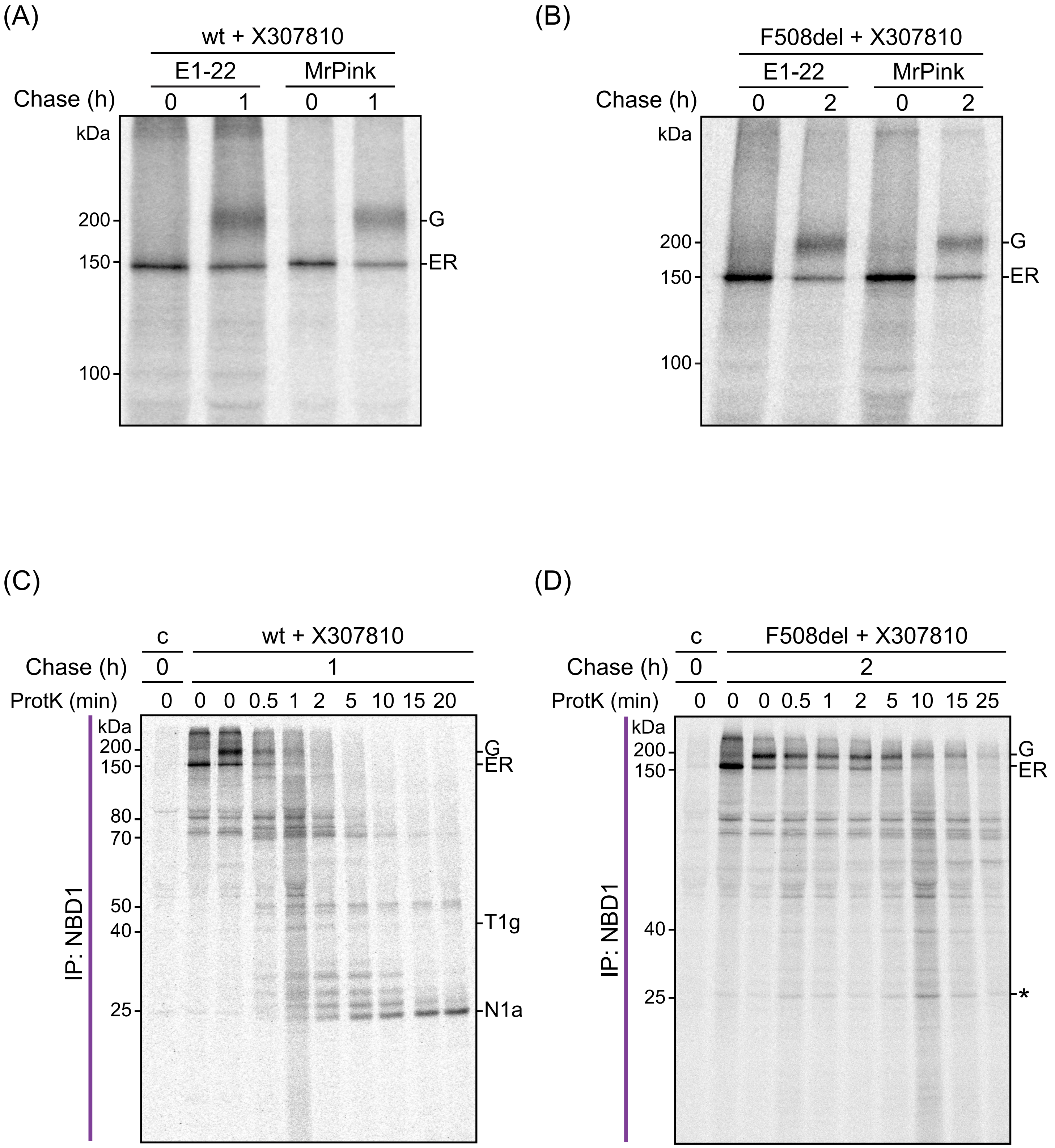
